# Accurate tracking of lytic granules via simultaneous particle tracking, phase retrieval and point spread function reconstruction

**DOI:** 10.1101/2025.05.02.651986

**Authors:** Mohamadreza Fazel, Reza Hoseini, Maryam Mahmoodi, Kevin L. Scrudders, Ayush Saurabh, Lance W. Q. Xu, Zeliha Kilic, Behshid Kasmaie, Mingdong Liu, Douglas Shepherd, Shalini T. Low-Nam, Fang Huang, Steve Pressé

## Abstract

3D tracking and localization of particles, typically fluorescently labeled biomolecules, provides a direct means of monitoring cellular transport and communication. However, sample-induced wavefront distortions of emitted fluorescent light as it passes through the sample and onto the detector often yield point spread function (PSF) aberrations, presenting an important challenge to 3D particle tracking using pre-calibrated PSFs. PSF calibration is typically performed outside cellular samples, ignoring sample-induced aberrations, which can result in localization errors on the order of tens to hundreds of nanometers, ultimately compromising sub-diffraction limited tracking. In practice, correcting sample-induced aberrations currently requires sample-specific hardware adjustments, such as adaptive optics. Yet, information on sample-induced aberrations and PSF shape can be directly decoded from data collected using a 3D imaging setup (*e*.*g*., bi-focal). To this end, we propose a framework for simultaneous particle tracking, phase retrieval, and PSF reconstruction (SPT-PR) directly from the input data themselves. We apply it to sub-diffraction tracking of lytic granules released at the immunological synapse of T cells revealing slower motions in proximity of the plasma cell membrane, consistent with assembly of the fusion machinery and, ultimately, degranulation and release of toxic payloads. To accomplish this, we operate within a Bayesian paradigm, placing continuous priors on all possible pupil phase and amplitudes warranted by the data without limiting ourselves to a finite Zernike set–thereby allowing capture of intricate pupil phase details. We benchmark our framework using a wide range of synthetic and experimental data from static to diffusing particles, and generalize to multiple diffusing particles with overlapping PSFs. Further, as a result of simultaneous particle tracking, phase retrieval, and PSF reconstruction, we retrieve the pupil phase with errors smaller than 10% under a range of realistic scenarios while demonstrating that for tracking lytic granules under an idealized Gaussian PSF assumption, we recover discrepancies as large as hundreds of nanometers.

## 1 Introduction

Fluorescence-based microscopy helps labeled biological samples standout from background [1,2]. Yet as fluorescent light, often emitted from labeled biomolecules of interest, propagates through the sample, the wavefront can be significantly distorted. These distortions arise from sample optical inhomogeneity, resulting in aberrated point spread functions (PSFs) [1, 3, 4]. These aberrations, in turn, hinder accurate 3D particle localization and tracking [5–7] using conventional methods relying on pre-calibrated PSFs or pre-assumed PSF shapes (*e*.*g*., Gaussian PSF) [2, 4, 8–13] (see Fig. 1 and S1).

**Figure 1:**
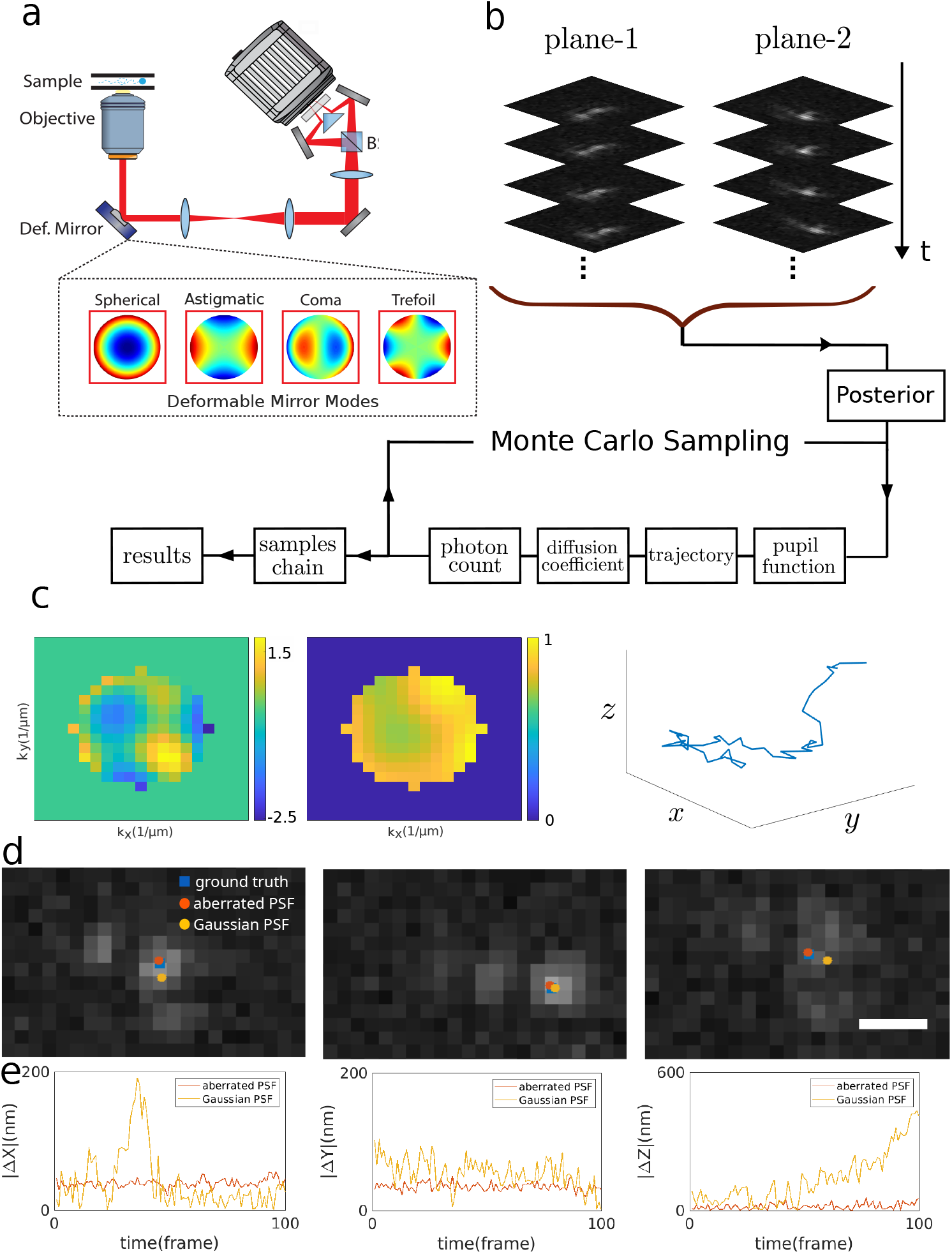
A depiction of a diffusing particle observed using a bi-focal microscope. The bi-focal setup provides two PSF slices separated by Δ*z* in the axial direction. A deformable mirror is used for wavefront manipulation. (a) Optical setup: fluorescent light emitted from a diffusing particle is collected by the objective and directed to the beam splitter (BS), resulting in two light beams with different path lengths. A deformable mirror in the light’s path is used here to induce known aberrations in the wavefront, serving as a test for the framework we propose; (b) computational framework: frame sequences from two planes are used to construct the probability (the posterior) over parameters of interest, namely, the pupil function, the particle’s trajectory, diffusion constant, and photon counts per frame. The framework we propose uses the entire sequence of bi-focal data to construct the posterior (the full joint probability over all possible particle tracks and the associated pupil phases and amplitudes). Probable values of the quantities of interest are then obtained by drawing samples from the posterior using Monte Carlo; (c) example of quantities of interest obtained: left to right panels, respectively, represent the most probable (maximum a posteriori) learned pupil phase, pupil amplitude and 3D trajectory; (d) example frames of a sequence of 100 frames of data simulated assuming a diffusing particle in the presence of optical aberrations with pupil phase and amplitude shown in panel c. Blue, red, and yellow markers, respectively, indicate true location, location found by our framework, and location found assuming a (best fit) Gaussian PSF. Scale bar is 500 nm; (e) differences of the true and recovered locations by our framework. Differences between the true location and those learned assuming a Gaussian PSF across all frames leads to localization errors, both laterally and axially, often exceeding hundreds of nanometers.

For instance, Fig. 1d-e illustrates frames of a distorted PSF arising from optical aberrations and the resulting localization errors induced by comparing the correct PSF (assuming known ground truth) to a commonly used Gaussian PSF idealization. Under this approximation, the resulting lateral and axial localizations differ from the ground truth by tens to hundreds of nanometers [14]; see Fig. 1e. As such, adaptive optics (AO) solutions are often used to correct for sample-induced wavefront distortions, bringing the PSF closer to an ideal Gaussian PSF [15–21].

Beyond PSF distortions, another major obstacle in 3D localization and tracking of labeled biomolecules arises from localization ambiguity in the axial direction arising from PSF symmetry with respect to the focal plane [7]. This issue is often addressed by using either multi-focal setups which break the symmetry along the axial direction [1, 22–25] or through PSF engineering which leverage designed wavefront perturbations [1, 26–33]

Yet, wavefront manipulation in both AO [15, 18, 19] and PSF engineering [1, 26–33] requires specialized optical hardware, such as cylindrical lenses, phase masks, deformable mirrors (DM) and spatial light modulators (SLM), each of which comes with its own limitations. For instance, SLMs are sensitive to light polarization, reducing the collected photon budget by half, and both DM and SLMs require specialized expertise to incorporate, calibrate, and control [2].

In addition to these experimental solutions to 3D localization and aberration correction, complementary computational approaches exist to address both challenges [3, 13, 34–41]. For instance, in 3D particle localization and tracking, the pupil function–a complex quantity describing the light’s wavefront and its changes due to sample-induced aberrations, instrument imperfections, PSF engineering, and others–is often obtained using phase retrieval techniques such as Gertchberg-Saxton (GS) [3, 34, 35], maximum likelihood [36, 39, 42], and deep learning [38, 40, 41]. These phase retrieval techniques rely on a stack of available calibration image frames acquired from static beads placed on the microscope stage and incrementing its location below and above the focal plane. The calibrated pupil function may then be used to generate a pre-calibrated PSF later employed in particle localization and tracking [5, 43]. However, because this phase retrieval procedure is conventionally performed using static beads outside the cellular environment, it is difficult to account for sample-induced aberrations [34–41].

Another class of phase retrieval techniques, such as INSPR [3] and uiPSF [13] rely on stochastic and axial sampling of the underlying 3D PSF using sparse blinking of static single molecules originating from an extended 3D biological structure. These techniques are therefore challenging to apply for cases where the sample is not static, such as in the case of tracking, 3D extent is not guaranteed, axial range is limited, and overlapping PSFs could be present.

Yet it is possible to avoid approximations such as pre-calibrated PSFs in tracking, and account for sample-induced aberrations–ultimately enabling 3D *in vivo* tracking with sub-diffraction resolution by achieving simultaneous phase retrieval and tracking; see Fig. 1. To this end, we introduce a Bayesian framework for simultaneous particle tracking and phase retrieval (SPT-PR) that operates directly on image-frame sequences. By treating phase retrieval, localization, and trajectory inference as a unified problem, the framework leverages information that is otherwise discarded when these tasks are performed in separate stages [2, 7]. In addition, photophysical parameters, such as intensity fluctuations, which are often ignored, are naturally incorporated and exploited to improve inference. This holistic approach therefore enables more accurate estimation of motion parameters, including particle trajectories and diffusion constants.

We initially benchmark the proposed framework using a wide range of both simulated data and *in vitro* data for a single static particle and a single diffusing particle, respectively. We show that our framework can accomplish sub-diffraction precision in both lateral and axial directions using data from molecules moving hundreds of nanometers below and above the focal plane, while simultaneously deducing the diffusion constants, and the pupil phase and amplitude due to optical aberrations with maximum errors of approximately 10% under typical imaging conditions. Next, we used SPT-PR to accurately track and study the trafficking of lytic granules in T cells during polarization. Delivery of lytic payloads to the plasma membrane is a rate-limiting factor in T cell cytotoxicity. T cells actively modulate this process to regulate the quantity of granules released at the immunological synapse. However, questions remain regarding how T cells modulate which granules are released and their dynamics [44, 45]. Here, using SPT-PR for tracking, we show that lytic granules slow down as they approach the plasma membrane, as expected [46]. We further compare these trajectories to those obtained using an idealized Gaussian PSF, demonstrating that Gaussian PSFs can introduce biases on the order of tens to hundreds of nanometers specifically in the axial direction.

## 2 Results

Our SPT-PR framework operates on a complete set of input frame sequences, denoted by 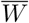 (with *W* representing a single frame), acquired from a multi-focal imaging system with *R* focal planes. The framework simultaneously and self-consistently infers the pupil amplitude *A*, pupil phase *ϕ*, 3D particle trajectory 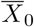, diffusion constant *D*, particle brightness (number of photons from particle per frame) *I*_*r*_, and uniform background *B*_*r*_ for each focal plane *r*. By simultaneously estimating the particle trajectories together with the pupil phase and amplitude, the method accounts for sample-induced aberrations, enabling accurate particle tracking deep within optically heterogeneous environments.

To simultaneously learn the unknowns, we operate within a Bayesian paradigm learning full distributions over all unknown parameters. As such, our results are often represented as histograms or medians of the resulting distributions for each parameter. Additionally, uncertainties ascribed to these parameters–drawn from the finiteness of data, detector features, and Poisson noise–are represented by the widths of the obtained distributions, indicating the spread of the posteriors corresponding to these parameters.

In what follows, we demonstrate our SPT-PR framework’s ability to retrieve pupil phase and amplitude from diffusing particles while simultaneously learning their trajectories with sub-diffraction limited precision, diffusion constant, and other parameters using a range of synthetic, *in vitro*, and *in vivo* data.

We benchmark our framework under various conditions, including different optical aberrations (pupil phases and amplitudes), diffusion constants, signal-to-noise ratios (SNR), and extend our framework to the multi-diffusing particle case with overlapping PSFs. Specifically: 1) we compare our SPT-PR framework’s performance in learning pupil phase and amplitude to conventional phase retrieval methods, such as GS algorithm [34, 35] and uiPSF [13], using static particle data (known particle locations) as, to our knowledge, SPT-PR is the only tool available for non-static particles. For this, we employed both simulated and *in vitro* data from static beads with various induced aberrations (see Figs. 2 and S2-S4); 2) we then highlight the novelty of our SPT-PR framework by extending beyond static samples in retrieving pupil phase and amplitude from a single diffusing particle while simultaneously learning the particle’s trajectory using both synthetic and *in vitro* data under different aberrations, diffusion constants, and SNR conditions (see Figs. 3 and S5-S7); 3) we benchmark SPT-PR by simulating a particularly challenging data set with multiple overlapping aberrated PSFs. SPT-PR uses this data set to simultaneously retrieve the pupil phase and amplitude and learn the particle trajectories along with the other parameters (see Fig. 4); finally, 4) we use SPR-PR to study the dynamics of lytic granules within T cell cytoplasm during polarization (see Fig. 5).

**Figure 2:**
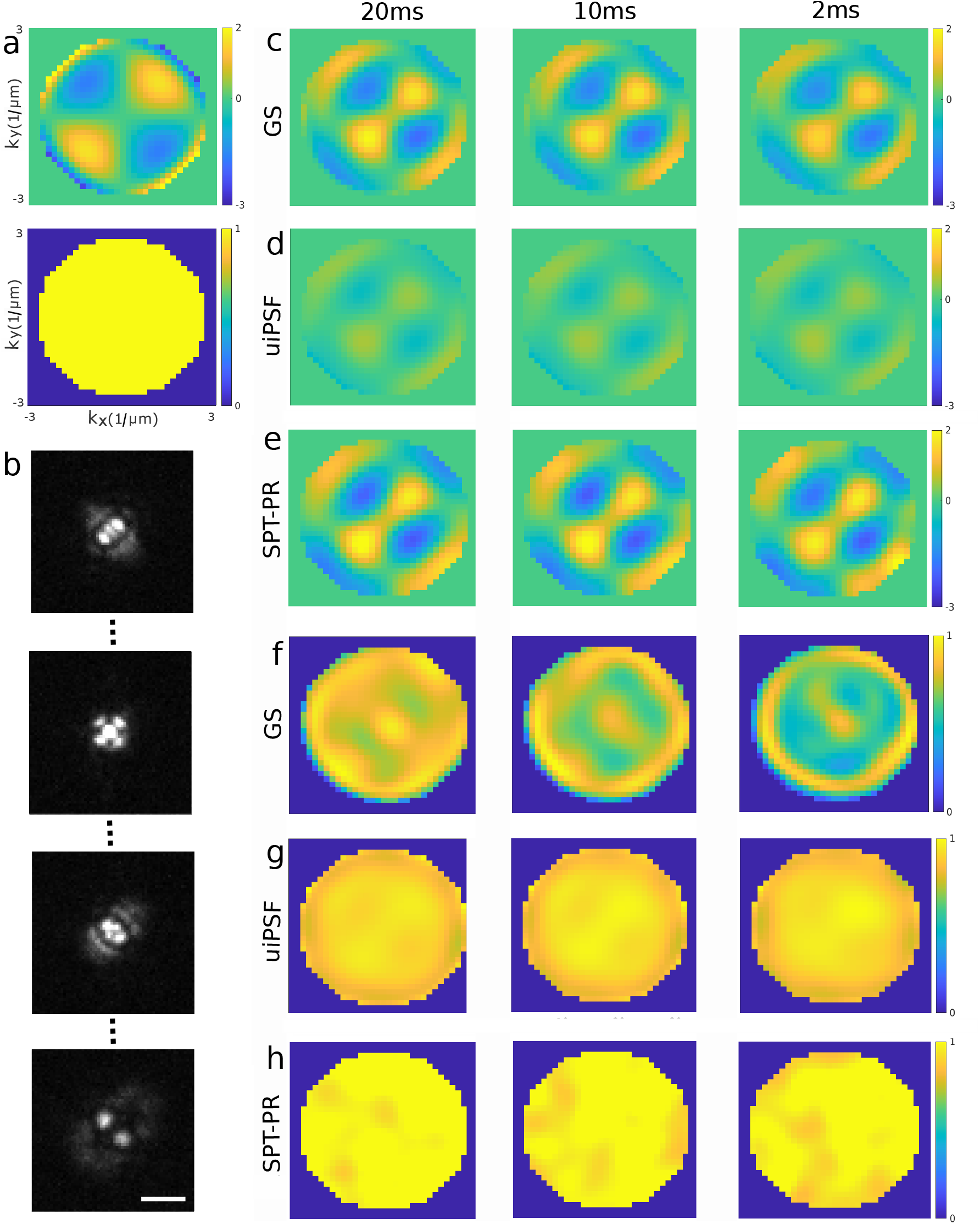
Phase retrieval for static beads using Gerchberg-Saxton, uiPSF, and SPT-PR. (a) ground truth induced phase (secondary astigmatic) and amplitude; (b) example PSFs at different Z-positions; (c) resulting phases from the GS algorithm using data collected with 20 ms, 10 ms, and 2 ms exposure times from left to right where by varying the exposure time we obtained different levels of SNR; (d) phases reconstructed using the resulting Zernike coefficients from uiPSF; (e) resulting phases from our SPT-PR framework; (f) resulting amplitudes from GS; (g) resulting amplitudes from uiPSF; (h) resulting amplitudes from our SPT-PR framework. The data was collected by placing a 100 nm bead on stage of a bi-focal microscope (data from only one of the planes was used for the analysis) and incrementing its axial location in 100 nm steps from -2 *µm* to 2 *µm*, numerical aperture (*N*_*a*_) of 1.35, refractive index of the medium in which the particle is embedded 1.449 (80% glycerol solution) and an emission wavelength of 680 nm, a 62×62 ROI with pixel size of 119 nm. The parameters are the same across the figures unless stated otherwise. The axes labels and ranges in panels c-h are similar to those in panel a. The colorbar unit is radian and this unit is retained throughout the manuscript. The scale bar is 2*µm*.

**Figure 3:**
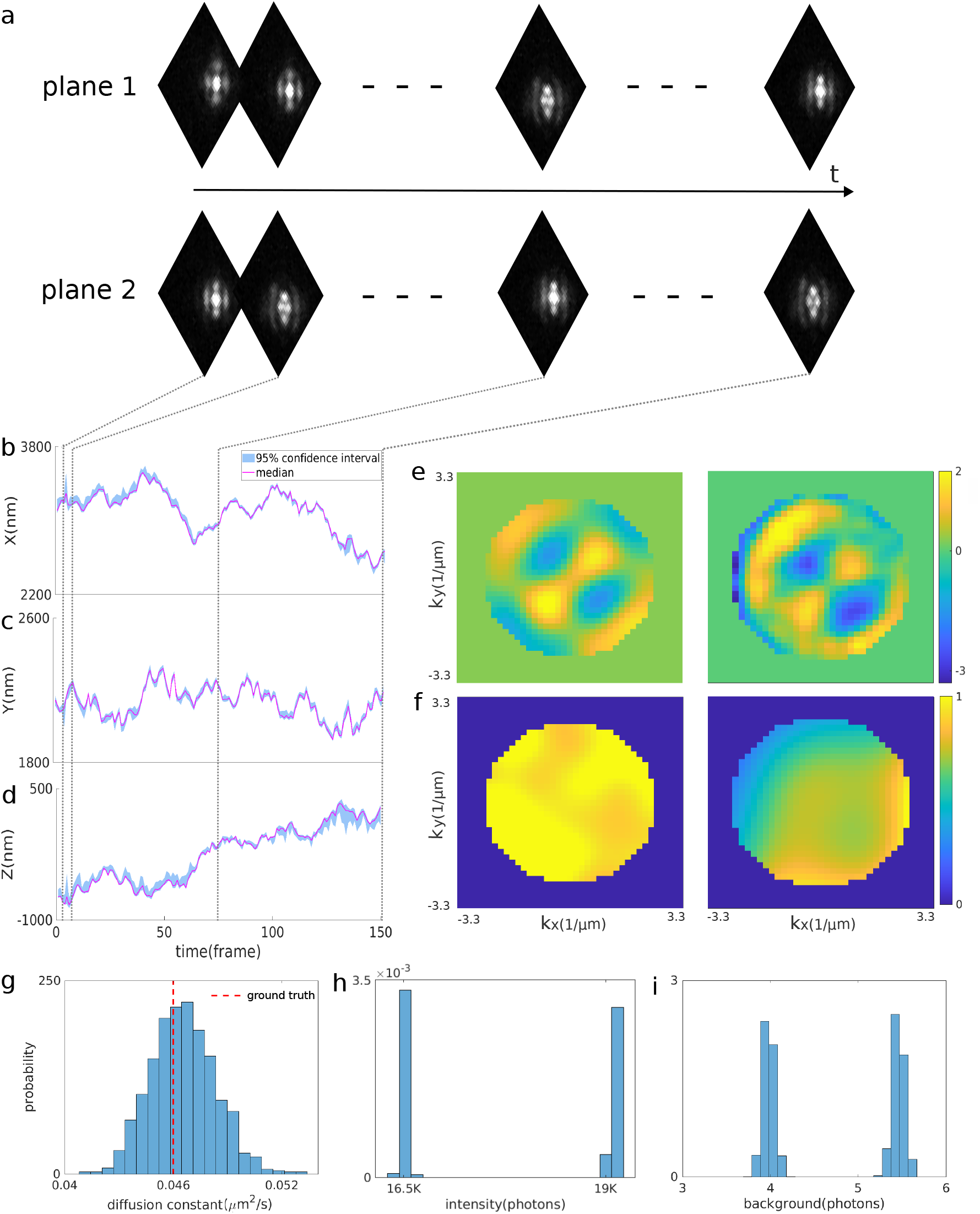
Simultaneous phase retrieval and particle tracking using *in vitro* data acquired from diffusing beads with 100 nm diameter within 80% glycerol solution, a camera exposure time of 20 ms, and induced secondary astigmatic aberration on top of aberrations due to refractive index mismatch. (a) examples of collected PSFs at different locations; (b) X-trajectory with average 95% credible interval of 22 nm; (c) Y-trajectory with average 95% credible interval of 23 nm; (d) Z-trajectory with average 95% credible interval of 43 nm; (e) from left to right, respectively, inferred pupil phases using static (see Fig. 2) and diffusing beads obtained using data sets acquired from the same setup with identical induced aberrations (secondary astigmatic); (f) from left to right, respectively, inferred pupil amplitudes using static and diffusing beads; (g) histogram of sampled diffusion constants. The red dashed line indicates the ground truth diffusion constant calculated using the Stokes-Einstein diffusion equation for 80% glycerol solution at room temperature; (h) the two histogram peaks show sampled particle intensities for two focal planes; (i) two histogram peaks representing sampled background for the two focal planes. All parameters are similar to Fig. 2, and we used 150 frames from each focal plane.

**Figure 4:**
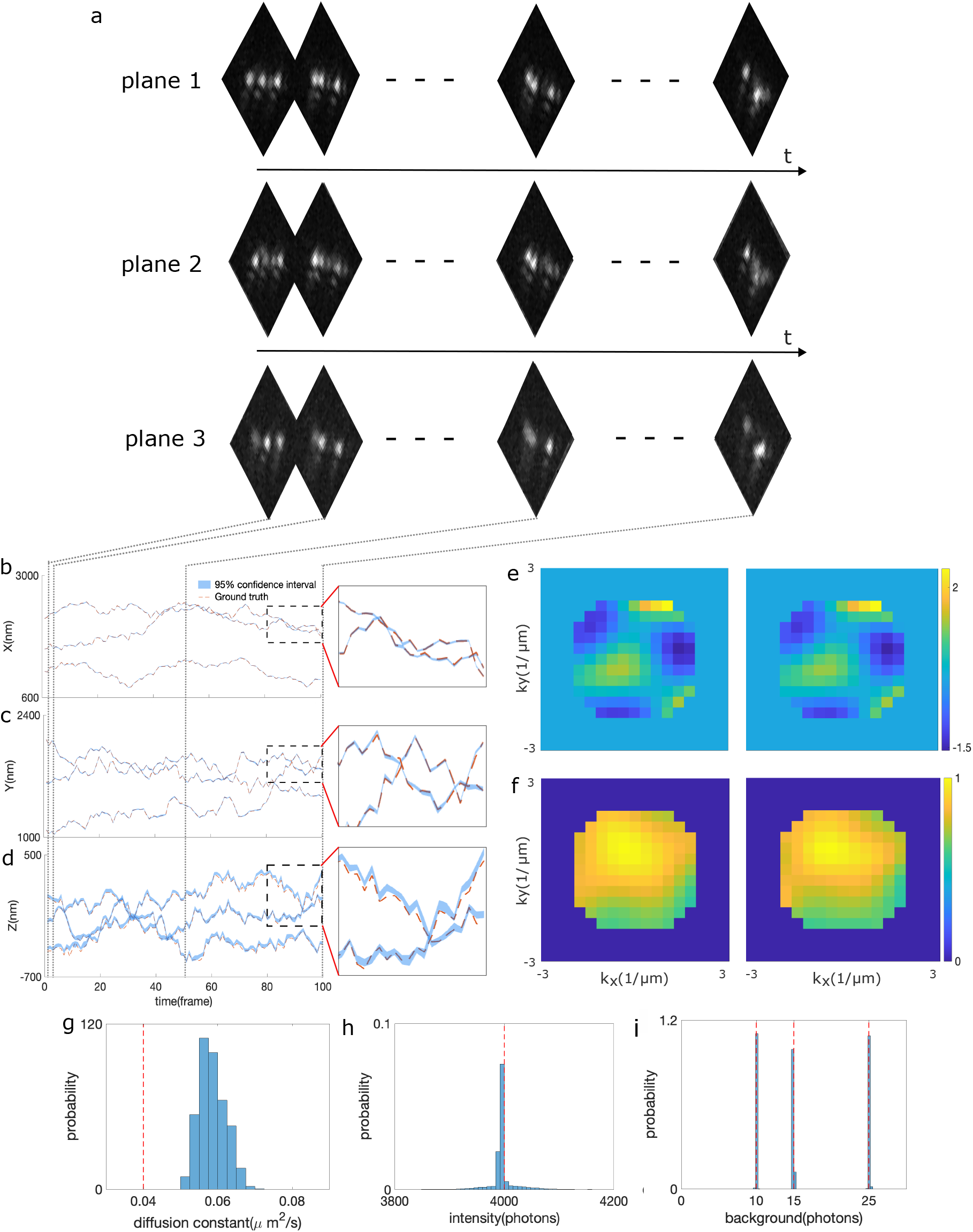
Simultaneous phase retrieval and particle tracking using synthetic data simulated assuming three particles with overlapping PSFs, with randomly generated pupil phase and amplitude, diffusion constant of 0.04 *µm*^2^*/s*, an intensity of 4000 photons per frame per particle, backgrounds of 10, 15, and 20 photons per pixel for different planes. (a) Examples of the simulated data assuming three focal planes; (b) X-trajectories; (c) Y-trajectories; (d) Z-trajectories; (e) the left and right panels, respectively, represent the ground truth and found pupil phases; (f) the left and right panels, respectively, represent the ground truth and found pupil amplitudes; (g) histogram of sampled diffusion constants. The diffusion constant histogram has a peak deviating from the ground truth due to lower localization accuracies in regions with overlapping PSFs; (h) histogram of sampled intensities (similar across focal planes and particles); (i) histograms of sampled backgrounds. All the other parameters used are similar to Fig. 2 except we assumed 100 frames of 32×32 pixels from each focal plane.

**Figure 5:**
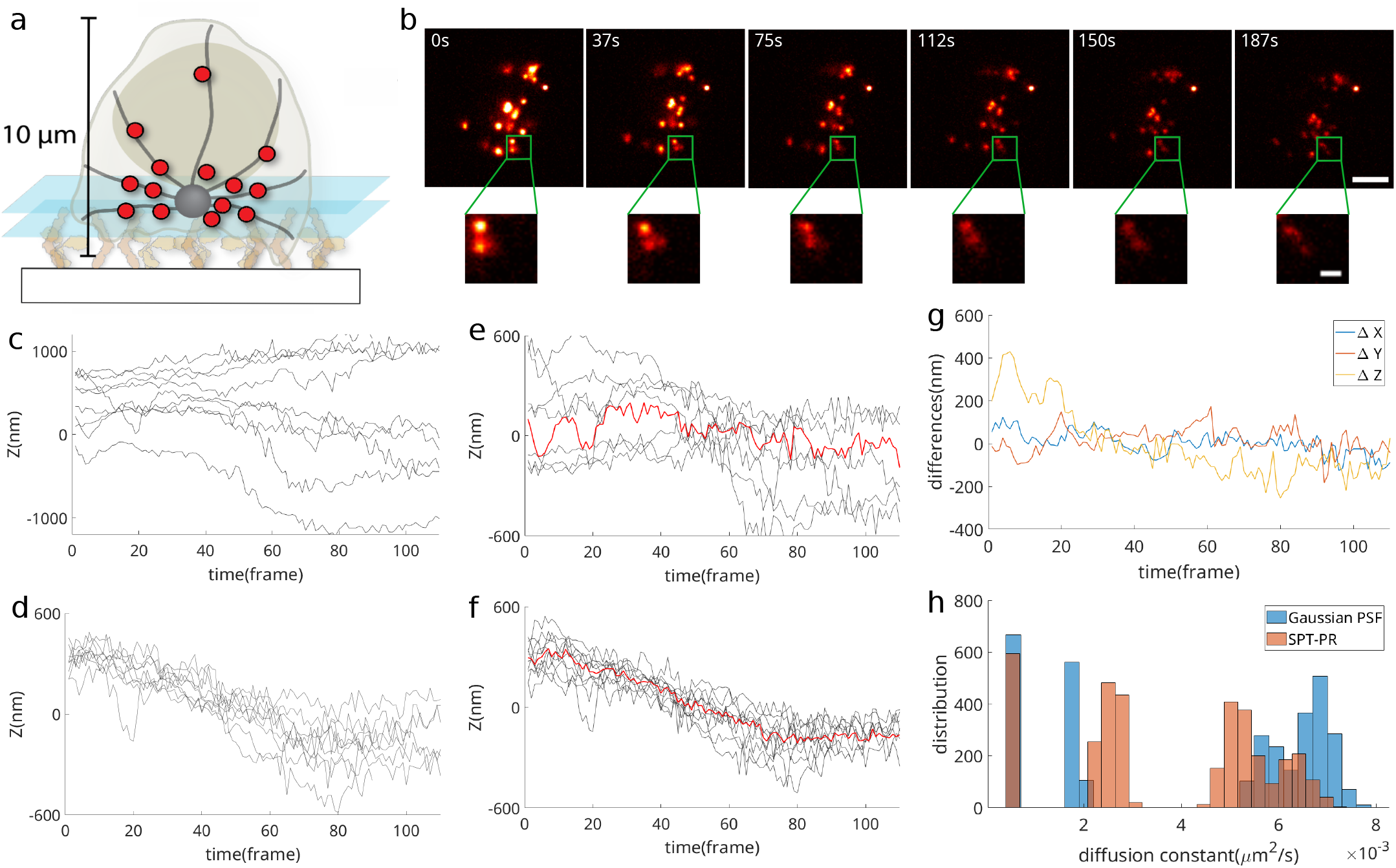
Tracking lytic granules in stimulated T cells while simultaneously learning the PSFs. (a) Schematic of T cells on activating substrate and polarizing cytotoxic granules (red circles) that traffic along microtubules (black lines). The white rectangle at the bottom indicates the glass coverslip and the blue planes are the foci of the bi-focal setup. (b) Example frames (different time points) of a data set acquired from granules loaded with LysoTracker Deep Red diffusing within Jurkat T cells. The indicated times are rounded to the nearest integer. Here, the frame exposure time and the time-lapse between the consecutive frames are, respectively, 100 ms and 1.6 s. The Green box indicates a selected ROI containing three overlapping PSFs used for granule tracking. Zoomed-in ROIs are shown at the bottom. Scale bars are, respectively, 4 *µm* and 1 *µm*. (c) The medians of the last 500 sampled Z-trajectories for 10 lytic granules in stimulated cells assuming an idealized Gaussian PSF. (d) The medians of the last 500 sampled Z-trajectories for 10 lytic granules using SPT-PR. (e) The Z-trajectories shown in panel c after subtracting the mean position of each trajectory prior to plotting, allowing trends in granule motion to be more clearly visualized. The red curve represents the median trajectory across the 10 granules. (f) The obtained Z-trajectories in panel d after subtracting the mean position of each trajectory. (g) Differences between the median trajectories obtained using an idealized Gaussian PSF and SPT-PR in the X, Y, and Z directions (e.g., the difference between the red curves in panels e and f for the Z-trajectories). (h) Histograms of sampled diffusion constants using an idealized Gaussian PSF and SPT-PR. The corresponding X and Y trajectories are provided in Fig. S11.

### 2.1 Phase retrieval using static particles

As an initial benchmark, we begin by demonstrating our SPT-PR framework in retrieving pupil phase and amplitude using a simple case: static beads. We compare our results with conventional phase retrieval methods: the standard GS algorithm [3, 34, 35] and Zernike fitting-based retrieval [42] as implemented in uiPSF [13]. To do so, we used both simulated and experimental *in vitro* data with a range of induced aberrations. The *in vitro* data were acquired from 100 nm (diameter) beads in an 80% glycerol solution (refractive index of 1.499) with a fluorescent emission wavelength of 680 nm using a bi-focal setup with a 370 nm inter-plane distance, and an effective pixel size of 119 nm. The fluorescent light was captured by an objective with a numerical aperture of 1.35, while the microscope stage was axially stepped from -2 *µm* to 2 *µm* in 100 nm increments. We used the deformable mirror implemented in this setup to obtain data sets with induced secondary astigmatic aberration (Fig. 2) and without any intentionally induced aberration (Figs. S2) and frame exposure times of 20 ms, 10 ms, and 2 ms. We also simulated data sets using similar parameters, both with and without aberrations (Figs. S3 and S4).

We first processed the data sets without intentionally induced aberrations (see Figs. S2, S3, and S8a). To do so, we cropped 100 frames of a 62×62 pixel region of interest (ROI) centered on the PSF from each plane for the analysis. As conventional phase retrieval methods assume known particle locations, for the sake of a head-to-head comparison of SPT-PR with conventional methods, we provided the particle location as input to our code and inferred the remaining parameters. Figs. S2 and S3 show the resulting pupil phases and amplitudes from SPT-PR, uiPSF, and GS, indicating agreement across all methods except the case with the lowest SNR, as anticipated. We next consider more complex cases by processing experimental data sets with induced aberration in a similar fashion to that described above. Fig. 2c-e, from left to right, shows the resulting pupil phases from GS, uiPSF, and our SPT-PR framework, using data sets with exposure times of 20 ms, 10 ms, and 2 ms, respectively, with approximately 30K, 10K, and 2K photon counts per frame on average, respectively. While the resulting phases from GS and our framework (Fig. 2c-d) cover the same range (colorbar from -3 to 2 radians), the resulting phase from uiPSF in Fig. 2d is underestimated. This underestimation is perhaps due to the constraint introduced in the uiPSF loss function to smooth the results [13]. In addition, Figs. 2f-h depict the resulting pupil amplitudes from GS, uiPSF, and our SPT-PR framework, respectively. The amplitude from GS is not flat as expected in the absence of attenuation [1] with approximately 8% and 21% error for data sets with 20 ms and 2 ms exposure times, and the uiPSF amplitude is under-estimated compared to those from GS and SPT-PR.

We further repeated the analyses with synthetic data where ground truth pupil phase and amplitude are available to quantify the errors in pupil phase and amplitude. For this purpose, we simulated data sets with optical aberrations using a combination of primary spherical and vertical trefoil aberrations (9th and 11th Zernike polynomials using the Noll index) for pupil phase, a flat pupil amplitude (Figs. S4), while other parameters were similar to those in the experimental data Fig. 2. The PSF corresponding to these aberrations is shown in Fig. S8b. Instead of varying exposure times, we assumed varying fluorescent emissions of 20K, 10K, and 2K photons per frame and a uniform background of 15 photons per frame over each pixel, motivated by the experimental data. The resulting phases and amplitudes are presented in Fig. S4c-h. Similar to the experimental results, we observed that uiPSF underestimates the phase, and the resulting phase from GS in Fig. S3 has an average of approximately 10% root mean square error in the estimated pupil phase.

### 2.2 Phase retrieval using a single diffusing bead

Here, we demonstrate the novelty of our SPT-PR framework in retrieving pupil phase and amplitude from a diffusing bead while simultaneously learning the bead’s trajectory alongside other parameters. This allows us to integrate the effect of sample-induced aberration, not otherwise currently achievable by other methods. To do so, we use *in vitro* and simulated data with a range of aberrations.

We begin by discussing our data and results for the *in vitro* data sets shown in Figs. 3 and S5. The data set in Fig. 3 was collected using a setup and sample similar to that described in section 2.1, though this time the stage remained static, but the bead was allowed to diffuse within an 80% glycerol solution, with an induced secondary astigmatic aberration on top of the aberration due to refractive index mismatch. A total of 2000 frames with an exposure time of 20 ms were collected from each plane. We selected a subset of the data by cropping a 62×62 pixels ROI from 150 consecutive frames (Fig. 3a).

The resulting bead trajectories are shown in Fig. 3b-d, where the cyan curves represent the median of the last 10K sample trajectories (drawn from the posterior using Monte Carlo), and the blue regions indicate the corresponding 95% credible intervals. In Fig. 3e-f, the left panels show the pupil phases and amplitudes obtained from a static bead (Fig. 2e and h), which we use as ground truths. The right panels depict the resulting pupil phase and amplitude from our SPT-PR framework, respectively. To assess the performance of SPT-PR in learning the pupil phase and amplitude from a diffusing bead, we calculated the average relative error between the ground truth and recovered values (given as (*ϕ*_true_ − *ϕ*_found_)*/*max(*ϕ*_true_)) in panels Fig. 3e-f to be approximately 26%. The relatively high error is due to the shift observed on the top left part of the found phase, while the shape of the phases from static and diffusing particles match with good accuracy. This shift is likely due to the existing errors in the microscope plane registration, which involved finding shifts and rotations across the two planes (see Supplementary Note 1). Moreover, uncertainty in the found pupil and amplitude is shown in Fig. S9. The constructed PSF from the found pupil function and the PSF uncertainties are also given in Fig. S8c and Fig. S10a. The histogram of the sampled diffusion constants is presented in Fig. 3g with the ground truth deviating from the histogram peak by less than 5%. Here, we used the Stokes-Einstein diffusion equation [47] to find the ground truth diffusion constant for 100 nm beads diffusing within an 80% glycerol solution at room temperature. Fig. 3h-i display histograms of the sampled background photons and particle intensities, each showing two peaks corresponding to the two planes in the optical setup.

We further tested our framework’s ability to learn the pupil phase and amplitude using two different frame subsets from the same *in vitro* data set of diffusing beads. The full data set consisted of 2000 frames, from which we cropped two 32×32 pixel ROIs over two separate 120-frame segments (Fig. S5a and d). We analyzed each segment independently and compared the resulting pupil phases and amplitudes, assuming that optical aberrations are approximately uniform over the field of view. Figs. S5b-c show the resulting pupil phases and amplitudes from two different segments of a data set with induced secondary astigmatic aberration plus aberration due to the mismatch between glycerol’s refractive index and the immersion oil. We calculated the average relative error between the phases and amplitudes, resulting in values of approximately 14% and 11%. Similarly, we selected two segments from another data set with induced coma and quadrafoil aberrations, repeating the analysis as described. The resulting pupil phases and amplitudes are shown in Figs. S5e-f. We also calculated the relative average error for pupil phases and amplitudes from these segments, finding values of approximately 12% and 4%, indicating good similarity between them.

After assessing our SPT-PR framework using experimental data, we tested our framework using simulated data with a single diffusing particle with available ground truth values. In contrast to experimental data, here, the available ground truth values allow for error quantification in the inferred quantities of interest. Therefore, we simulated a data set with parameters similar to our experimental conditions: a diffusion constant of 0.04 *µm*^2^*/s*, a media refractive index of 1.499, and an emission wavelength of 680 nm. We also assumed fluorescence emission collected using a bi-focal setup, with an inter-plane distance of 370 nm, a numerical aperture of 1.35, a pixel size of 100 nm, and 100 frames of 32×32 pixels from each plane.

Fig. S6a shows example frames of simulated data generated using the specified parameters and a randomly generated pupil phase and amplitude (a superposition of multiple Zernike polynomials; Fig. S6e-f, left panels). The PSF corresponding to these aberrations is shown in Fig. S8d. The learned trajectory is shown in Fig. S6b-d, where the ground truth trajectory (dashed lines) lies within the 95% credible interval of the sampled trajectory (blue area) for all three coordinates. The retrieved pupil phase and amplitude are shown in the right panels of Fig. S6e-f, with less than 2% average relative error compared to the ground truth values on the left panels. Histograms of the sampled diffusion constants, particle intensity, and background photon counts are shown in Fig. S6g-i, respectively, with the ground truth values (dashed red lines) within the histogram ranges and deviations from the peaks under 3%.

Further, we simulated data sets with parameters similar to those in Fig. S6 but varied diffusion constants and particle brightness ranging from 0.03-0.09 *µm*^2^*/s* and 500-4000 photons, respectively (see Fig. S7). Fig. S7 depicts histograms of the sampled diffusion constants for these cases. As expected, the histograms sharpen with increasing particle brightness. However, as the diffusion constants increase, the peaks of the histograms begin to deviate from the ground truth values. This occurs due to the motion blur effect distorting the PSF shape for faster diffusing particles, leading to larger errors in reconstructed PSFs and, consequently, lower precisions in the learned diffusion constants.

### 2.3 Phase retrieval using multiple diffusing particles with overlapping PSFs

In this section, we challenge our SPT-PR framework with data sets simulated assuming three particles with overlapping PSFs (Fig. 4). Here, we simulated data from three focal planes with different background levels of 10, 15, and 25 photons per pixel per frame, and ∓370 nm separation along the axial direction (Fig. 4a). Additionally, we used 4000 photons per frame per particle and a randomly generated pupil phase and amplitude (Fig. 4e-f, left panels), while keeping all other parameters similar to those in Fig. 3. The PSF corresponding to this pupil function is shown in Fig. S8e.

Our SPT-PR framework learned the correct trajectories shown in Fig. 4b-d. The ground truth trajectories (dashed lines) lie within the 95% credible intervals of the sampled trajectories (blue regions). The deduced pupil phase and amplitude, displayed in the right panels of Fig. 4e-f, deviate from the ground truth values (left panels) by less than 4% using the average relative error defined earlier. Histograms of the sampled diffusion constants, particle intensity, and background are shown in Figs. 4g-i, respectively.

While the ground truth values for intensity and background (red dashed lines) fall within the histogram ranges, the peak of the diffusion constant histogram deviates from the ground truth by approximately 30%. This is due to reduced localization accuracies (approximately 21 nm and 40 nm on average in the lateral and axial directions, respectively) and larger areas where the ground truth trajectories happen to be slightly outside of the 95% credible interval (for instance see Fig. 4d), particularly in regions with heavily overlapping PSFs.

### 2.4 Lytic granules tracking using SPT-PR and idealized Gaussian PSFs

We used our SPT-PR framework to track and study the diffusion of lytic granules during T cell polarization in response to activating surfaces (Fig. 5a). The resulting trajectories were then compared with those obtained using conventional Gaussian PSF–based tracking approaches (Fig. 5 and S11). Cells were imaged using a bi-focal setup with 370 nm separation in the axial direction, 100 ms exposure time and a time lapse of 1.6 s between consecutive frames. Cells were caught proximal to landing on the surface to ensure early detection of polarization, within the first few minutes of interaction with the substrate. Figure 5b shows representative frames of the acquired data from these diffusing granules. Tracking was performed on regions of interest (ROIs) containing one or more granules whose motions spanned a few hundred nanometers below and above the focal planes while remained within the field of view for approximately 100 consecutive frames. This enabled accurate extraction of trajectories, pupil phases and amplitudes, PSFs, and other relevant parameters. Figure 5c–d show the inferred axial trajectories of 10 granules, while the corresponding lateral trajectories are presented in Fig. S11. To better visualize the directional trends in axial motion, the mean position of each trajectory was subtracted in Fig. 5e–f.

Figure 5g shows the differences between the median trajectories of the 10 granules obtained using Gaussian PSFs and SPT-PR across different spatial directions. These differences range from tens to hundreds of nanometers, with the largest discrepancies observed along the axial direction, reaching up to 400 nm. This highlights the strong sensitivity of Z-localization to accurate PSF modeling. Consistent with this difference, the trajectories obtained using an idealized Gaussian PSF model (Fig. 5e) do not exhibit a clear directional bias, whereas those obtained with SPT-PR (Fig. 5f) reveal a pronounced tendency toward lower axial (*Z*) positions. Such axial errors in granule trajectories are significant and may promote misinterpretations of the underlying phenomena. In the case of granule polarization, some granules might be interpreted as hovering rather than being fully transported toward the intermembrane interface. Even if hovering occurs, a location of 50 nm or 200 nm above the plasma membrane is critical since close approximation is necessary for assembly of the fusion machinery and release of the cytotoxic contents. By comparison, SPT-PR shows the robust binarization of the cellular decision to polarize and demonstrates coordination between the intracellular machinery to transport the granules. This is an interesting realization given the range of sizes of granules.

Finally, Figure 5h displays histograms of the average diffusion constants over full granule trajectories for each ROI. However, the obtained trajectories shown in Fig. 5f indicate a much slower dynamics along the axial direction near the end of their motion as they approach the plasma membrane. To quantitatively evaluate this observation, we directly calculated diffusion constants using only the last 30 localizations of each obtained trajectory (see Supplementary Section 3), corresponding to the stage at which the lytic granules are in close proximity to the plasma membrane. This analysis resulted in average values of 1.2 × 10^−3^*µm*^2^*/s*, consistently lower than those obtained for the full trajectories indicating that granules diffusion slows as they approach the membrane. The corresponding retrieved pupil phases and amplitudes are shown in Figs. S12 and S13.

## 3 Theoretical Methods

Here, we provide a brief description of our framework. We first briefly introduce the likelihood of a single focal plane and then extend it to the case of multi-focal microscopes [3]. Further details of the framework can be found in the Supplementary Information.

In the far field limit, the PSF is related to the pupil function, 𝒫(*k*_*x*_, *k*_*y*_), via scalar diffraction theory [1, 7, 35]

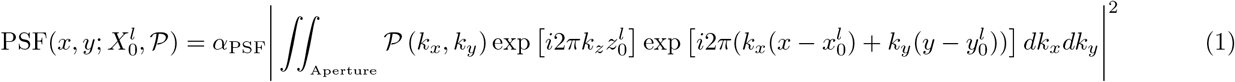

where *α*_PSF_, 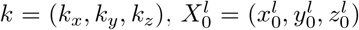, and (*x, y*), respectively, denote the normalization constant, the wave vector, particle coordinates at the *l*th frame (time point), and the camera coordinates. Further, the pupil function is a complex quantity expressed by its phase and amplitude *𝒫* = *Ae*^*i*Φ^. The axial component of the wave vector is given by 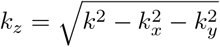. The integral is performed over the range of wave vectors accessible through the microscope aperture (objective) given by 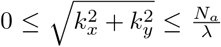 where *N*_*a*_ and *λ* are, respectively, the numerical aperture and fluorescent light wavelength in vacuum. The above integral is approximated using sums computed employing fast Fourier transforms; for details, see Supplementary Information Section 1.

Assuming Brownian motion, the transition probabilities for particles from frame to frame, then given the particle location at frame *l*, the location at frame *l* + 1 follows

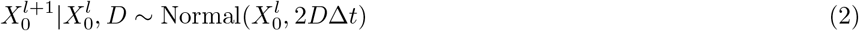

where *D* and Δ*t*, respectively, denote the variance in step size (also termed diffusion constant in the language of diffusion processes) and frame exposure time.

Using the resulting PSF eq. (1), the expected number of photons reaching the *n*th pixel of the *l*th frame then follows from [7, 48, 49]

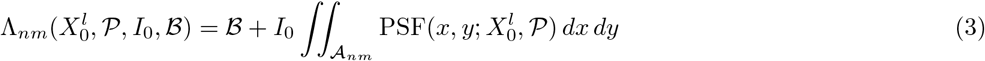

where *I*_0_ and *ℬ* are, respectively, the total photon count (intensity) from the particle per frame and the background photons per pixel. Here, *𝒜*_*nm*_ represents pixel area on the *n*th row and *m*th column of a 2D frame. Eq. (3) can be extended to the case with multiple particle by summing over the photon contribution from each particle; see SI Section 1. We express 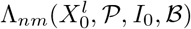 as 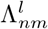 from hereon for notational simplicity. Given the expected photon counts in eq. (3), the likelihood of the photon count *ζ*_*nm*_ for the pixel in the *n*th row and *m*th column follows a Poisson distribution. However, the reported pixel values (data designated by 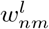) are not the direct photon counts. The reported pixel values are further contaminated by noise due to stochastic photon count conversion into pixel values in cameras. The camera noise follows a Normal distribution, leading to the following likelihood for the entire frame sequence [1, 50]

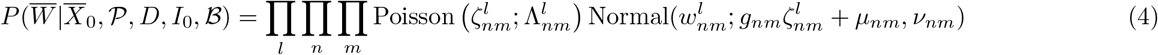

where 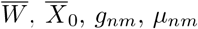 and *ν*_*nm*_, respectively, denote the entire set of frame sequences, the particle’s trajectory across a given sequence of frames, pixel-wise camera gain, offset and noise variance. These camera parameters can be calibrated in advance and are assumed to be known.

The likelihood obtained can be used to learn the set of unknown parameters. However, this likelihood is often degenerate with respect to the particle trajectory due to the symmetric PSF shape below and above the focal plane [1]. That is, this likelihood does not result in a unique trajectory. To overcome this issue, we therefore use data acquired by a multi-focal setup which simultaneously samples the PSF at multiple *z*-locations with known separations. Therefore, the task of particle localization at a time point reduces to learning the particle’s coordinates in only one of the planes (reference plane) using data from all the planes, which in turn breaks the symmetry and allows us to learn the true underlying trajectory.

As such, we generalize the above likelihood to accommodate a multi-focal (often bi-focal) setup providing data from *R* PSF slices, designated by *r* = 1, …, *R*. The likelihood of eq. (4) then reads as follows

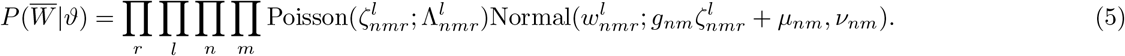

However, this likelihood model is computationally expensive, and thus, we use its approximate form provided in the Supplementary Information Section 1.

The likelihood obtained in eq. (5) can now be employed to estimate: the particle trajectory 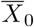, pupil function *𝒫* = *Ae*^*i*Φ^, diffusion constant *D*, particle photon emission rate (photon counts per frame *I*_*r*_ in plane *r*), and the background photon count per pixel *ℬ*_*r*_ in plane *r*. We collectively regroup these parameters under 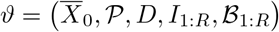.

We now proceed to construct the posterior by including priors on the unknown parameters

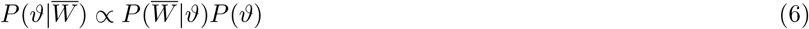

where *P* (*ϑ*) stands for priors on these parameters. Here, the most notable prior is the Gaussian process (GP) prior [51–55] on the pupil phase

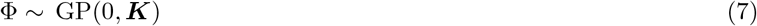

and similarly for the pupil amplitude we also use a GP prior; see Supplementary Information Section 2. Furthermore, the prior on the particle’s trajectory is given by eq. (2). For the rest of the parameters, we use priors that are either computationally or physically motivated; see Supplementary Information Section 2.

With the posterior at hand, we are now in a position to draw samples of the unknown parameters from the posterior. As the resulting posterior cannot be directly sampled on account of its non-analytic form, we use Markov chain Monte Carlo [53, 56–61] to iteratively sweep over the parameters. At each iteration, we sample the parameters in the following order: 1) sample the pupil function by drawing values from the phase and amplitude priors of corresponding pixels in Fourier space using Metropolis-Hastings (MH) [56,57]; 2) the particle’s trajectory is sampled by proposing new trajectories, *i*.*e*., set of particle locations across all the frames within a given sequence using a hit-and-run sampler [62,63]. That is, a new trajectory is proposed by displacing the particle from its current locations along random directions for each time point (frame); 3) sample step size variance by directly sampling from the posterior; 4) sample photon counts from the particle per frame using MH; 5) sample background per pixel again using MH. At the end, the set of samples drawn can be employed for further numerical analysis.

## 4 Experimental Methods

### 4.1 In-vitro fluorescent beads sample

High-precision glass coverslips (25 mm diameter, CG15XH, Thorlabs) were cleaned by sequential sonication for 15 minutes each in ethanol (2701, Decon) and HPLC-grade water (W5-4, Fisher Chemical), followed by drying with compressed air. Dark Red fluorescent beads (100 nm diameter, Invitrogen) were diluted 1:100,000 in deionized water.

1. *Static bead samples*: To prepare stationary bead samples, 200 *µL* of poly-L-lysine solution (P4707, Sigma-Aldrich) was applied to a coverslip and incubated at room temperature (RT) for 20 minutes. This was followed by incubation with 200 *µL* of the diluted bead solution for an additional 20 minutes. The coverslip was then rinsed with deionized water. For index-matched samples, 20 *µL* of 38% 2,2-thiodiethanol (TDE, 166782, Sigma-Aldrich) in 1×PBS (10010023, Gibco) was added. For index-mismatched samples, 80% glycerol was used instead. In both cases, a second cleaned coverslip was gently placed on top to form a sandwich structure.
2. *Diffusing bead samples*: To prepare samples with diffusing beads in an index-mismatched environment, 120 *µL* of the diluted bead suspension in 80% glycerol was deposited onto a cleaned coverslip, and another cleaned coverslip was placed on top.

For both stationary and diffusing bead samples, the coverslip sandwich was secured in a circular imaging chamber and sealed with two-component silicone dental glue (Twinsil Speed 22, Dental-Produktions und Vertriebs GmbH).

### 4.2 Live cell sample

Glass coverslips (25 mm diameter, 64-0735, Warner Instruments) were cleaned in a Hellmanex III solution (Z805939, SigmaAldrich) by sonication at 50°C for 30 minutes. Post-cleaning and rinsing in ultrapure water, coverslips were coated with poly-D-lysine (10 *µg*/mL, A38904-01, Gibco) and anti-CD3 (clone OKT3, 10 *µg*/mL, 566685, BD Biosciences) for 60 minutes at room temperature with gentle rocking. Jurkat T cells (generously provided by Dr. Arthur Weiss) were maintained in RPMI 1640 medium (11875093, ThermoFisher) supplemented with 10% v/v fetal bovine serum (FBS) and penicillin/streptomycin.

For live-cell imaging, cells were stained with LysoTracker Deep Red (100 nM in PBS, L12492, ThermoFisher) at 37°C for 10 minutes to label low-pH granules, including lytic granules. Excess LysoTracker dyes was not washed out, as it exhibits minimal fluorescence in neutral pH and becomes fluorescent upon accumulation in acidic granules. Following staining, 200 *µL* of the cell suspension in Live Cell Imaging Solution (supplemented with 1% m/v BSA, and 6 mM D-glucose, A59688DJ, ThermoFisher) was added onto a pre-warmed, coated coverslip, which was secured in an imaging chamber (Attofluor cell chamber, A7816, ThermoFisher). Cells were allowed to settle and establish contact with the functionalized surface for 10 minutes at room temperature. Cell interface establishment was monitored using a bright-field microscope. Simultaneous imaging was performed using a custom-built bi-focal super-resolution microscope to track the movement of lytic granules towards the coverslip interface and their diffusion within the cytosol.

### 4.3 Super resolution microscopy setup

The custom-built super-resolution microscope is based on a commercial inverted stand (IX-73, Olympus) and equipped with a 100× silicone oil immersion objective lens (NA 1.35, UPLSAPO100XS, Olympus America) mounted on a three-axis piezoelectric nano-positioning system (Nano-LP200, Mad City Labs). The system comprises two main optical pathways: illumination and emission. The illumination path incorporates three modules: wide-field excitation, guide-star generation, and bright-field illumination. Both wide-field excitation and guide-star generation utilize a 642 nm laser line (2RU-VFL-P-2000-642-B1R, MPB Communications), power-modulated via an acousto-optic tunable filter (AOTFnC-400.650-TN, AA Opto-electronic) to enable precise control of laser intensity. The emission path includes a primary detection path, containing a deformable mirror (DM), and a wavefront sensing path. Emitted fluorescence from the sample passes through a bandpass emission filter (FF01-731/137-25, Semrock) and is reflected by a deformable mirror (Multi-3.5, Boston Micromachines) positioned at the conjugate pupil plane. Fluorescence signals are then directed onto a 90° specialty mirror (47-005, Edmund Optics), generating a bifocal image plane, which is projected onto a scientific CMOS (sCMOS) camera. The effective pixel size at the sample plane is 119 nm, optimized for single-molecule localization and tracking.

## 5 Discussion

3D single particle tracking within cellular environments has been employed to study cell signaling [64], viscosity of cell cytoplasm during necrosis [65], and the diffusion of proteins involved in the organization of meiotic chromosomes [66]. However, accurate 3D particle tracking within cellular environments is often challenged by poor 3D particle localization due to sample-induced PSF aberrations. Moreover, current particle localization and tracking methods rely on PSF pre-calibrations [17, 67, 68]. Here, PSF calibration is often performed *in vitro* ignoring sample-induced aberrations and only considering aberrations from the optical setup. To address this critical issue in a self-consistent manner, here, we adopted a Bayesian framework to simultaneously deduce the sampled-induced aberrations (pupil phase and amplitude), reconstruct the PSF, and learn particle trajectories from the input data itself.

We benchmarked our SPT-PR framework using a range of synthetic and experimental data from both static and diffusing beads. First, to compare our framework to the conventional phase retrieval methods, such as GS and uiPSF, we used simulated and experimental data from static beads with and without optical aberrations. While uiPSF tended to under-estimate the phases, and the resulting amplitudes from GS deviated from the ground truths with decreasing photon counts, our framework generated consistent pupil phases and amplitudes across various photon counts (Fig. 2, Figs. S2-S4). Additionally, we used both simulated, *in vitro* experimental data from diffusing particles to demonstrate the novelty of our framework in retrieving the pupil phase and amplitude while learning our particle’s trajectory. Our Bayesian framework achieved sub-diffraction limited localization precisions with 95% credible intervals for the trajectories ranging from 12 nm to 24 nm in the lateral dimension and 21 nm to 48 nm in the axial dimension (Figs. 3, 4, and S6). Moreover, our framework deduces pupil phase using photon counts as low as 500 photons (Figs. S3, S4 and S7). What is more, we further challenged our framework using simulated data with multiple diffusing particles and overlapping aberrated PSFs. The resulting pupil phase and amplitude from our SPT-PR framework matched the ground truth with less than 5% error. Moreover, since our SPT-PR framework is capable of tracking particles deep inside the sample with high accuracies, we employed it to study the dynamics of lytic granules and observed that these granules slow down in the vicinity of plasma membrane while idealizations induced by assuming Gaussian PSFs struggled to show evidence of migration to the plasma membrane due to large axial errors, see Figs. 5 and S11.

Due to the sequential nature of our Monte Carlo sampling procedure, the computational cost is linearly dependent on the size of the frames, the number of frames, and the number of focal planes of input data. Furthermore, the computational cost is also tied to the nature of the optical aberrations. That is, optical aberrations with more intricate pupil phases require a larger number of samples to capture details of the pupil phase, leading to more computational complexity. For instance, it took about 1 hour to learn the phase shown in Fig. S6 while it took approximately 5 hours to learn the phase shown in Fig. 3 using a desktop with AMD Ryzen 9 3900X 3.8 GHz CPU and 64 GB RAM.

Also, while we have not yet fully exploited parallelism, our framework presents significant opportunities for optimization using modern computational hardware, particularly GPUs. Many of the computationally intensive components in our model can be naturally decomposed into independent calculations, making them well-suited for parallel execution. For instance, the integral in eq. (3) is evaluated independently for each pixel, allowing the workload to be distributed across multiple GPU cores and executed simultaneously. Leveraging such parallelism has the potential to dramatically accelerate computation and further enhance the scalability of our approach.

Although to reduce the computational time, we assumed a point-like fluorescent particle, the presented framework can be modified to accommodate extended light sources by convolving the extended particle shape (often a sphere) with the fluorescent emission in eq. (1). Moreover, the algorithm can be further modified to incorporate the motion blur effect by splitting the frame exposure time into multiple sub-periods and averaging over the resulting PSFs for each sub-period. The SPT-PR framework can also be extended to the nonparametric regime to deduce the number of particles within the input data, along with the other parameters. Further, accurate estimations of the pupil phase, amplitude and subsequently the PSF require that particle motion in the axial direction spans a few hundred nanometers above and below the focal plane, which may limit precise tracking of particles with restricted axial movement. This challenge can be mitigated by simultaneously analyzing multiple particles, each sampling a smaller axial interval. Finally, while we assumed an identical diffusion model for each particle, the provided framework can be generalized to accommodate an independent diffusion model (different diffusion constants or non-Brownian diffusions) for each particle.

## 6 Contributions

MF and SP conceived of the project. MF developed the model and the mathematical framework with help from ZK on the early formulation. MF and RH developed the software package. MF performed the analyses with help from RH, LWQX, AS, and BK. FH, MM, KLS, ML, and STL provided the experimental data. FH, AS, KLS, STL and DS provided helpful discussions. SP supervised and oversaw the entire project. MF wrote the paper with help from SP. All authors reviewed and approved the manuscript.

## 7 Code availability

The SPT-PR software package is available on Github: [SPT-PR software Package](https://github.com/MohamadFazel/Simultaneous-particle-tracking-phase-retrieval-and-PSF-reconstruction).

## 8 Data availability

Some of the data used in this study is included in the software package ([SPT-PR software Package](https://github.com/MohamadFazel/Simultaneous-particle-tracking-phase-retrieval-and-PSF-reconstruction)), and the rest is available upon request from the corresponding author.

## 10 Acknowledgment

SP acknowledges support from the NIH MIRA award (NIGMS R35GM148237). FH acknowledges support from the US National Institute of General Medical Sciences (R35GM119785). KLS was supported by Grant Number T32 GM132024 from NIH NIGMS. We thank Joerg Enderlein for useful discussions and comments, and Jay Spendlove for testing the software package.

## Supplementary Information

**Fig. S1:**
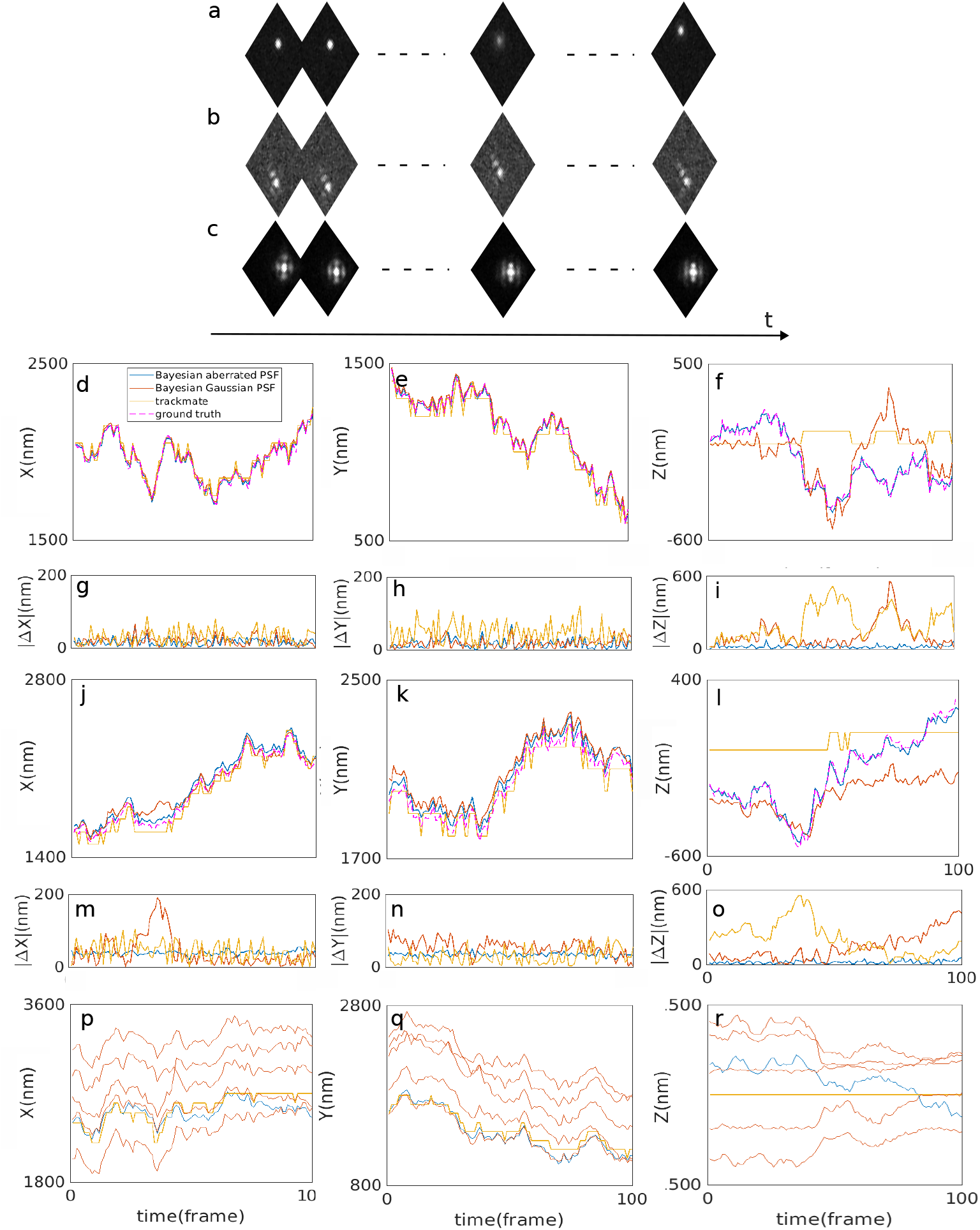
3D tracking using aberrated PSFs V.S. Gaussian PSF. Here, we compare resulting tracks using our method to simultaneously deduce trajectories and PSF shapes, and trajectories using Gaussian PSFs in a Bayesian framework [1] and Track-Mate [2]. (a) data simulated assuming a diffusing particle with non-aberrated (Gaussian) PSF; (b) data simulated assuming a diffusing particle with aberrated PSF; (c) experimental data acquired from a particle diffusing within a glycerol solution with an aberrated PSF; (d) X-trajectories from panel a; (e) Y-trajectories from panel a; (f) Z-trajectory from panel a; (g) difference of the ground truth X-trajectory from the resulting trajectories for panel a; (h) difference of the ground truth Y-trajectory from the resulting trajectories for panel a; (i) difference of the ground truth Z-trajectory from the resulting trajectories for panel a; (j) X-trajectories from panel b; (k) Y-trajectories from panel b; (l) Z-trajectories from panel b; (m) difference of the ground truth X-trajectory from the resulting trajectories for panel j; (n) difference of the ground truth Y-trajectory from the resulting trajectories for panel k; (o) difference of the ground truth Z-trajectory and the resulting trajectories for panel l; (p) X-trajectories from panel c; (q) Y-trajectories from panel c; (r) Z-trajectories from panel c. In panels p-r, although, we could not calculate the differences due to the lack of ground truth trajectories for experimental data, we note that more than one particle (multiple trajectories) was predicted when modeling an aberrated PSF using a Gaussian PSF. All the parameters for both simulated and real data are similar to Fig. 2 except we used 100 frames of data from a single focal plane, and a mirror reflection of the trajectory with respect to the focal plane was used when predicting on wrong Z-side for the Gaussian PSF due to its symmetry with respect to the focal plane.

**Fig. S2:**
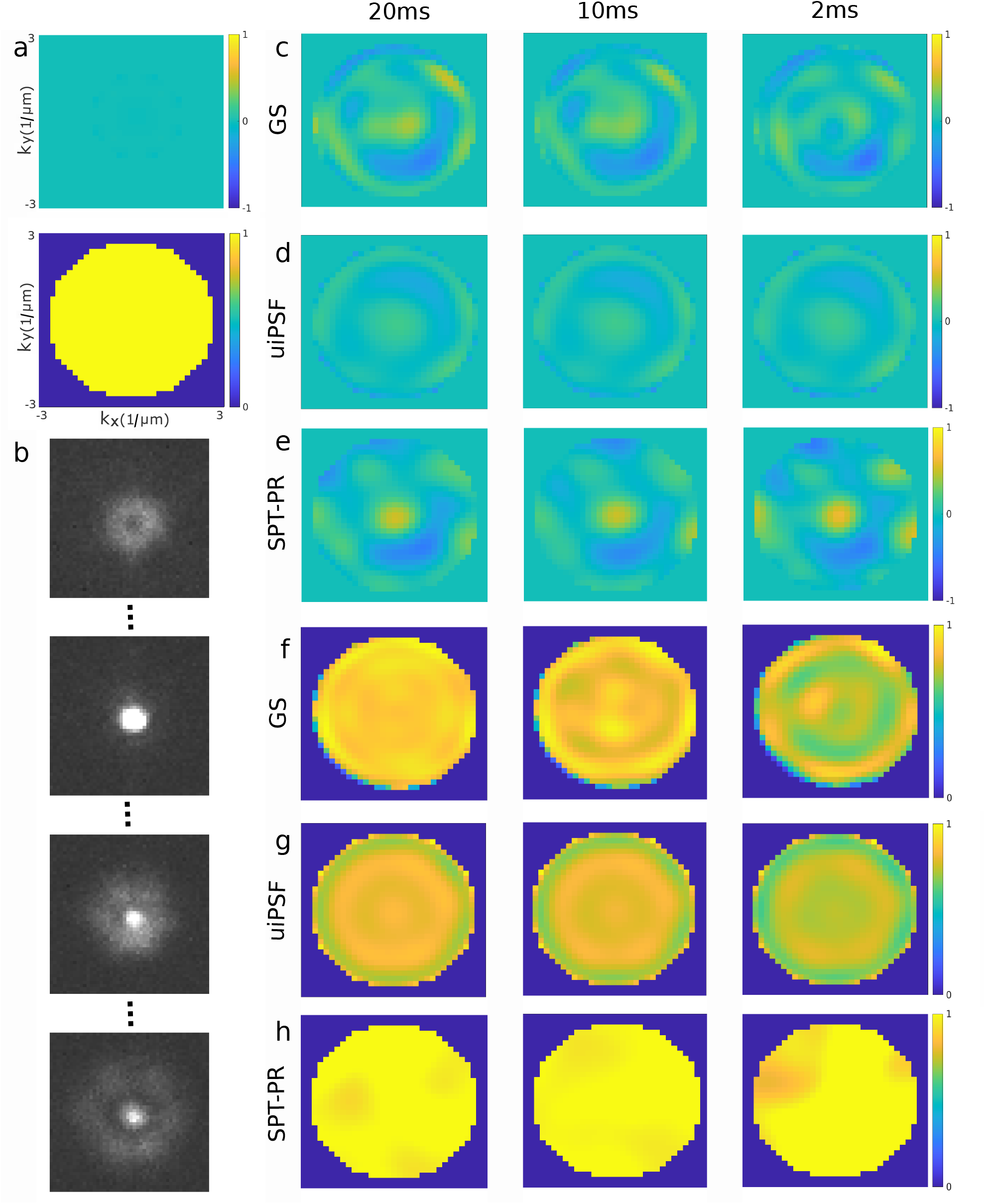
Phase retrieval for static beads using experimental data with no induced aberration using Gerchberg-Saxton, uiPSF, and SPT-PR methods. (a) ground truth phase and amplitude; (b) example PSFs at different Z-positions; (c) resulting phases from GS algorithm using data acquired with exposure times of 20 ms, 10 ms and 2 ms from left to right; (d) resulting phases from uiPSF; (e) resulting phases from SPT-PR; (f) resulting amplitudes from GS; (g) resulting amplitudes from uiPSF; (h) resulting amplitudes from SPT-PR. All the parameters are similar to Fig. 2. The axes labels and ranges in panels c-h are similar to panel a. The pupil phase and amplitudes from GS and uiPSF were reconstructed using their output Zernike coefficients in this figure, Fig. 2 and Fig. S3-4.

**Fig. S3:**
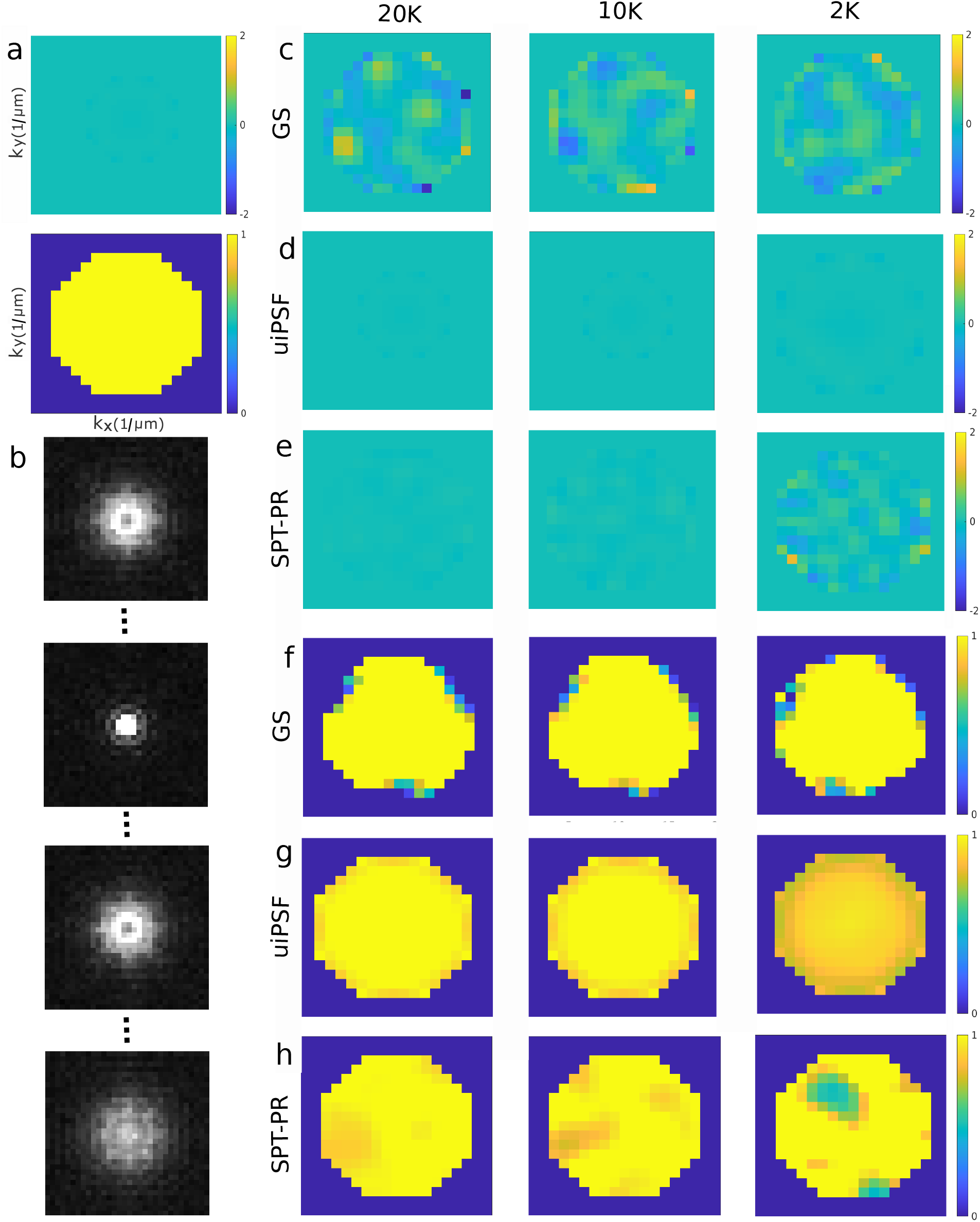
Phase retrieval for static beads using simulated data with no induced aberrations using Gerchberg-Saxton, uiPSF, and SPT-PR methods. (a) ground truth phase and amplitude; (b) example PSFs with no aberration at different Z-positions; (c) resulting phases from GS algorithm using simulated data by 30K, 10K and 2K photons from left to right; (d) resulting phases from uiPSF; (e) resulting phases from SPT-PR; (f) resulting amplitudes from GS; (g) resulting amplitudes from uiPSF; (h) resulting amplitudes from SPT-PR. To simulate the data sets used here, we assumed a bead is placed on a single plane microscope stage and its axial location is incremented in 100 nm steps from -2*µm* to 2*µm*. All the parameters are assumed to be similar Fig. 2, except we used ROIs with the size of 32×32 pixels. The axes labels and ranges in panels c-h are similar to panel a.

**Fig. S4:**
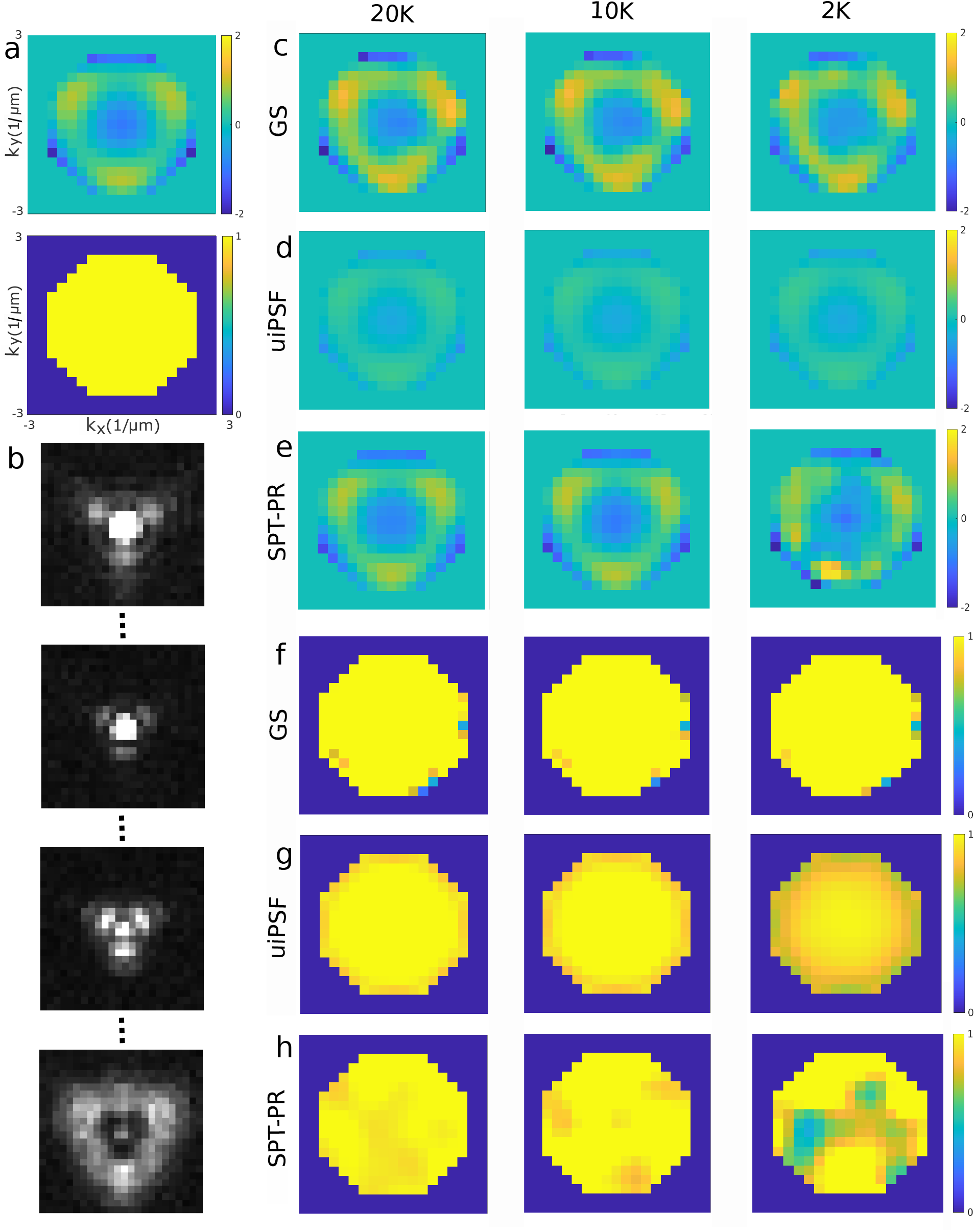
Phase retrieval for static beads using simulated data with induced aberrations using Gerchberg-Saxton, uiPSF, and SPT-PR methods. (a) ground truth phase and amplitude; (b) example PSFs at different Z-positions; (c) resulting phases from GS algorithm using simulated data by 30K, 10K and 2K photons from left to right; (d) resulting phases from uiPSF; (e) resulting phases from SPT-PR; (f) resulting amplitudes from GS; (g) resulting amplitudes from uiPSF; (h) resulting amplitudes from SPT-PR. The data simulation procedure is similar to Fig. S3. The axes labels and ranges in panels c-h are similar to panel a.

**Fig. S5:**
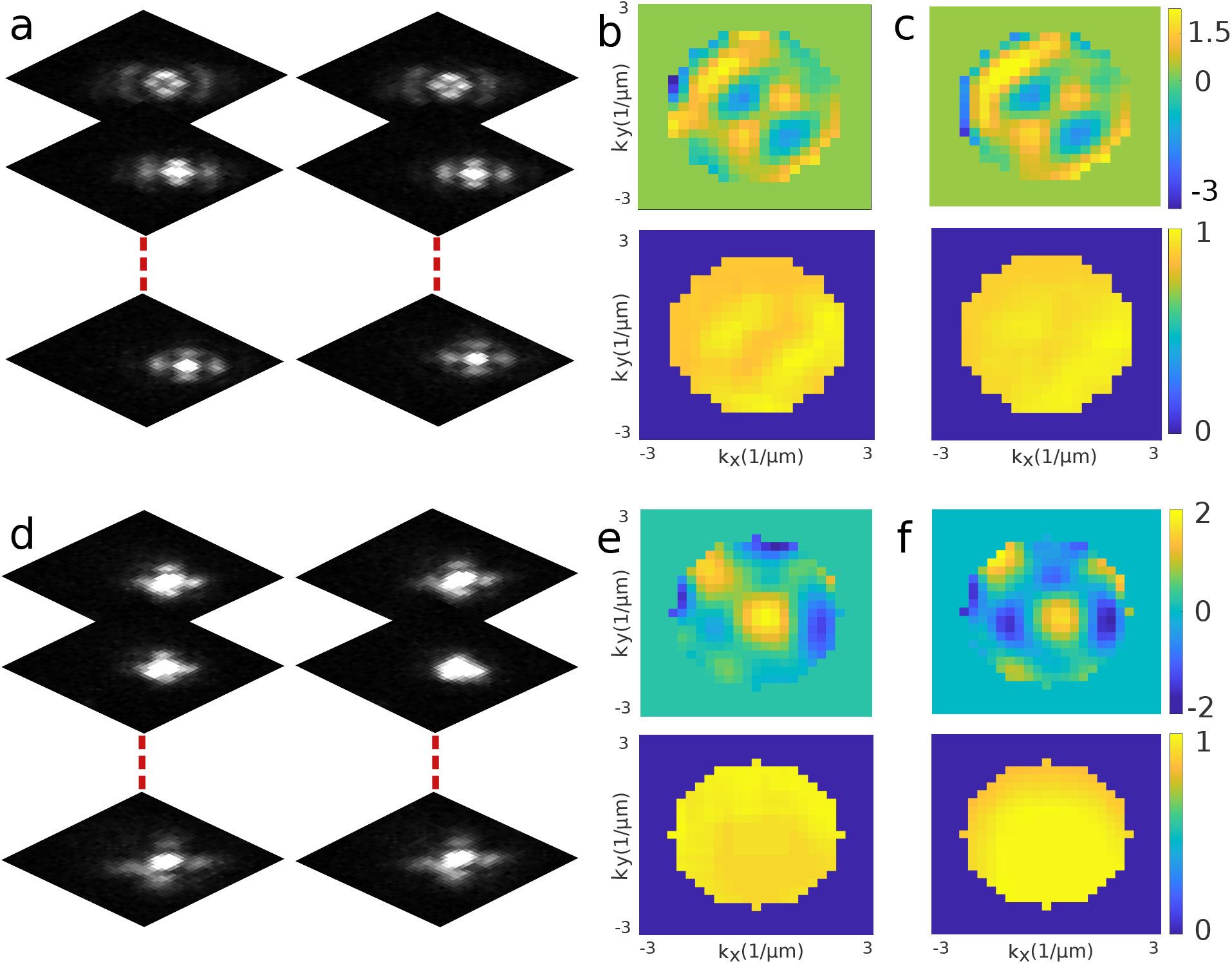
Simultaneous phase retrieval and particle tracking using *in vitro* data acquired from a bi-focal setup using 100 nm beads diffusing within 80% glycerol solution, a camera exposure time of 20 ms and various induced aberrations. (a) an example of frame sequences acquired using a bi-focal setup with secondary astigmatic induced aberration; (b) the top and bottom panels, respectively, show the pupil phase and amplitude obtained from a sequence of 120 frames of the data set with secondary astigmatic induced aberration; (c) the top and bottom panels, respectively, show the pupil phase and amplitude obtained from a different sequence of 120 frames selected from the same data set with secondary astigmatic induced aberration; (d) an example of frame sequences acquired using a bi-focal setup with three different induced aberrations including: primary astigmatic, Coma and Quadrafoil; (e) the top and bottom panels, respectively, show the pupil phase and amplitude obtained from a sequence of 120 frames of the data set with primary astigmatic, Coma and Quadrafoil aberrations; (f) the top and bottom panels, respectively, show the pupil phase and amplitude obtained from a different sequence of 120 frames selected from the same data set with primary astigmatic, Coma and Quadrafoil aberrations. All the parameters are similar to Fig. 3, except we used 32×32 pixel ROIs.

**Fig. S6:**
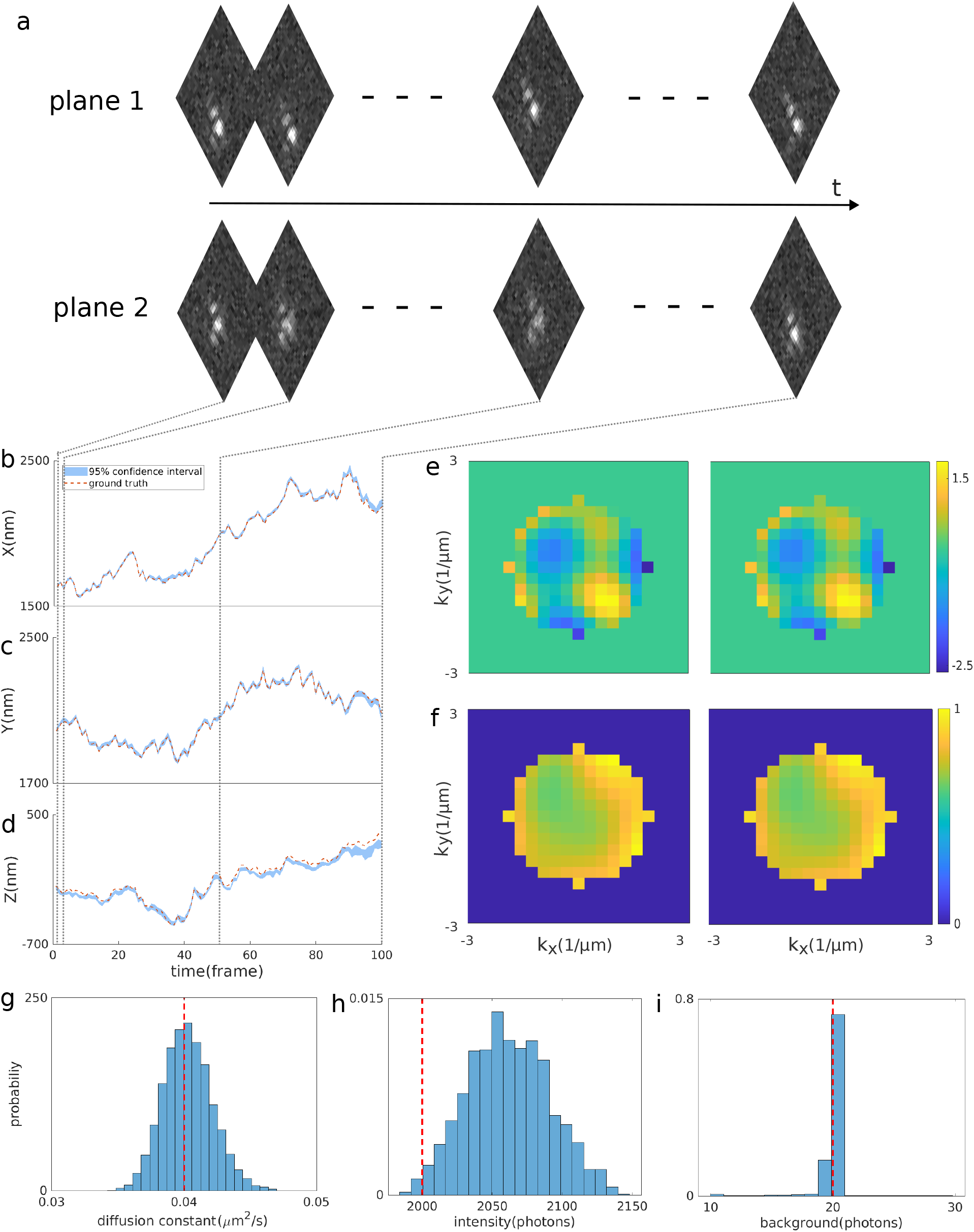
Simultaneous phase retrieval and particle tracking using synthetic data. (a) examples of simulated PSFs at different locations; (b) X-trajectory with average 95% confidence interval of 12nm; (c) Y-trajectory with average 95% confidence interval of 14nm; (d) Z-trajectory with average 95% confidence interval of 21nm; (e) the left and right panels, respectively, represent the ground truth and found pupil phase; (f) the left and right panels, respectively, represent the ground truth and found pupil amplitudes; (g) histogram of sampled diffusion coefficients; (h) histogram of sampled particle intensities; (i) histogram of sampled background. Red dashed lines indicate ground truths. All the parameters used in the simulations are similar to Fig. 3.

**Fig. S7:**
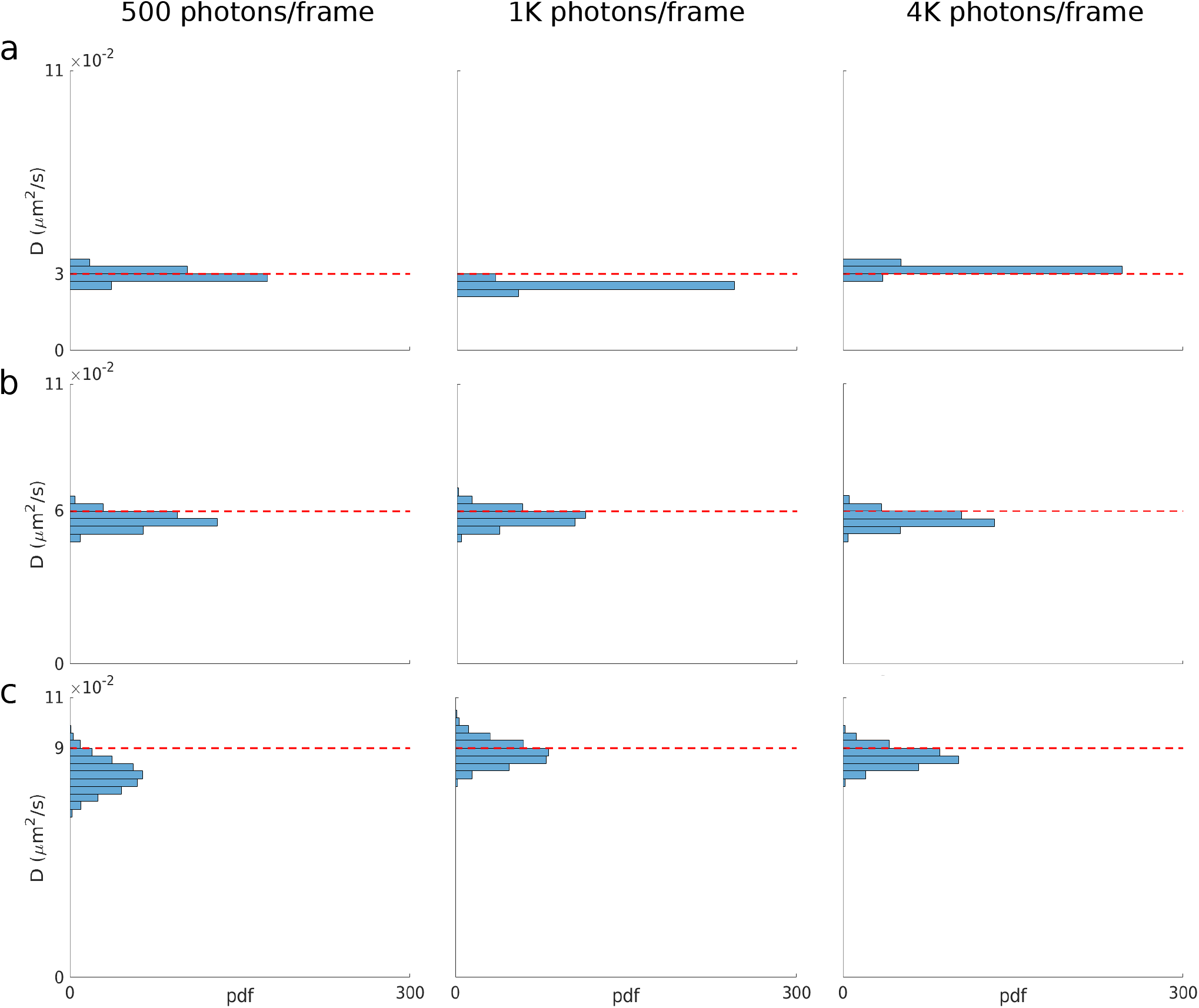
Control simulations–algorithm performance with increasing diffusion constant (D) and signal to noise ratio (SNR). Data sets were simulated using parameters (including pupil phase and amplitude) similar to Fig. S6 except we varied diffusion constant and particle brightnesses over a range of 3 × 10^−2^-9 × 10^−2^*µm*^2^*/s* and 500-4000 photons per frame per particle, respectively.

**Fig. S8:**
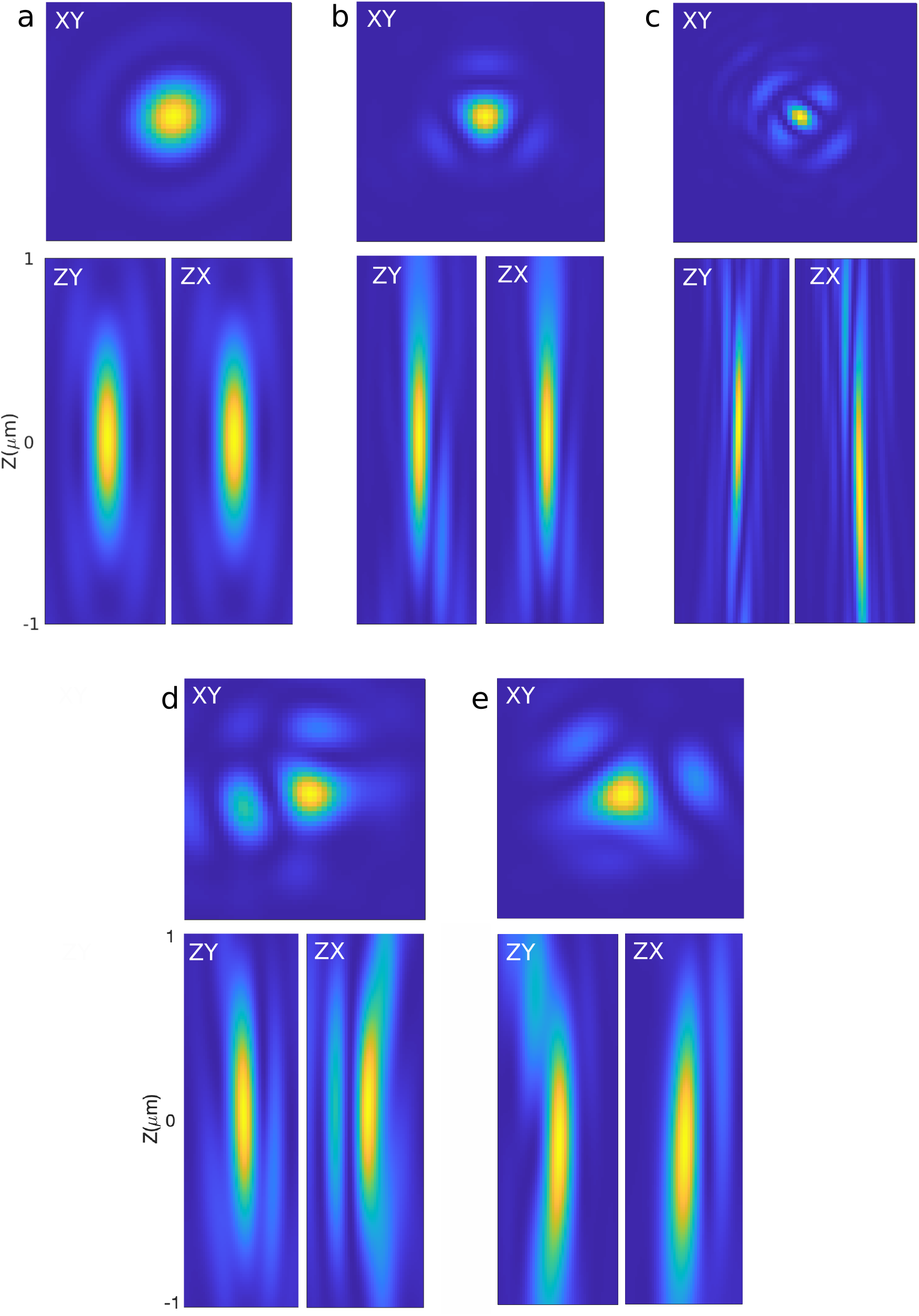
Different PSF cross-sections at *Z* = 0 (focus), *X* = 0 and *Y* = 0. (a) PSF corresponding to pupil function in Fig. S2-S3 (no aberration); (b) PSF corresponding to the pupil function in Fig. S4 (pupil phase simulated with 9th and 11th Zernike polynomial); (c) PSF corresponding to the pupil function in Fig. 2-3 (induced secondary astigmatism); (d) PSF corresponding to the pupil function in Fig. S6 (randomly generated pupil function); (e) PSF corresponding to the pupil function in Fig. 4 (randomly generated pupil function).

**Fig. S9:**
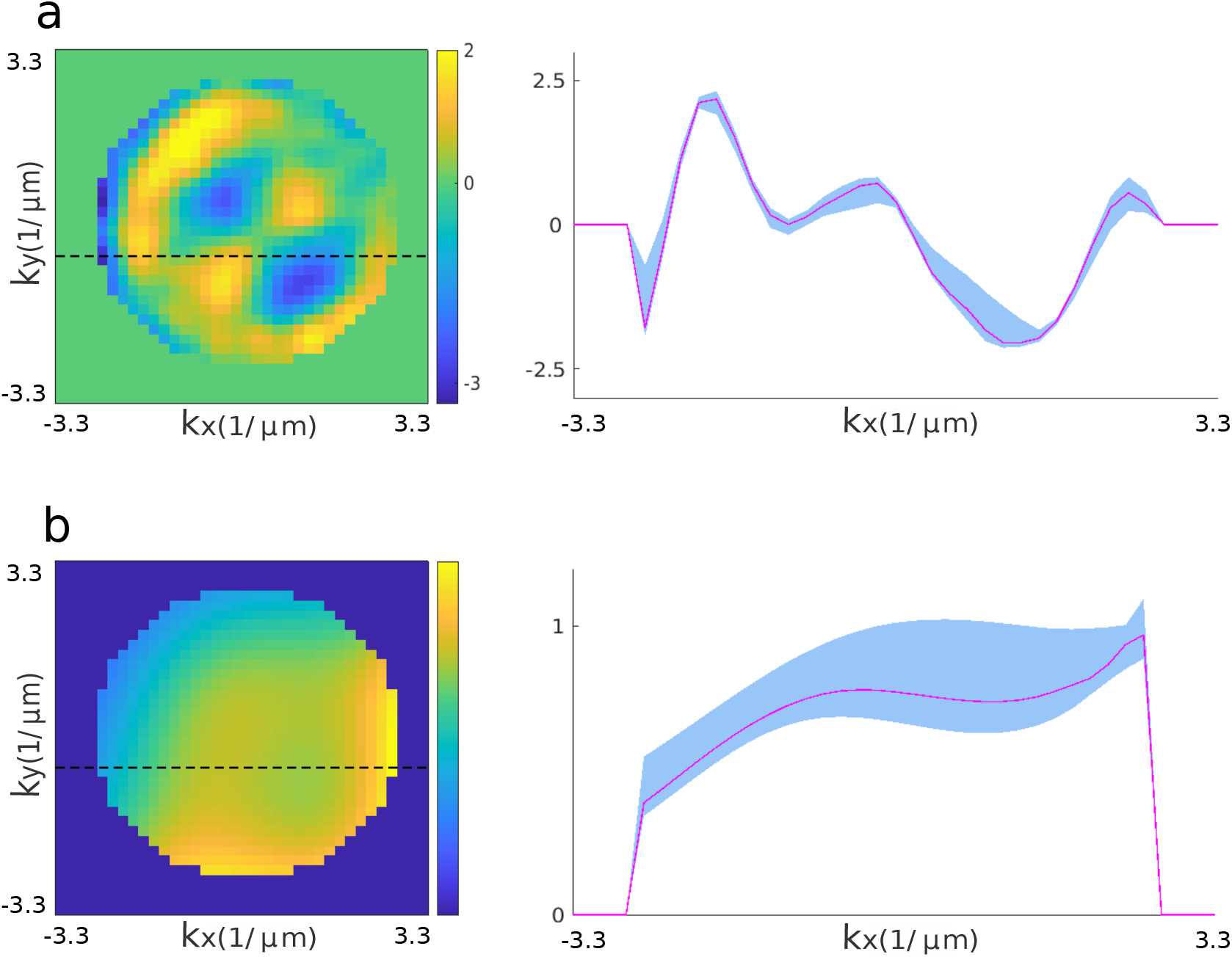
Uncertainties in the found pupil phases and amplitudes. (a) left and right panels, respectively, show the pupil phase in Fig. 3 and the uncertainty in the pupil phase estimation along the dashed line in the left panel; (b) left and right panels, respectively, show the pupil phase in Fig. 3 and the uncertainty in the pupil amplitude estimation along the dashed line in the left panel. Here, blue regions and cyan curves, respectively, designate the corresponding 95% confidence intervals and medians.

**Fig. S10:**
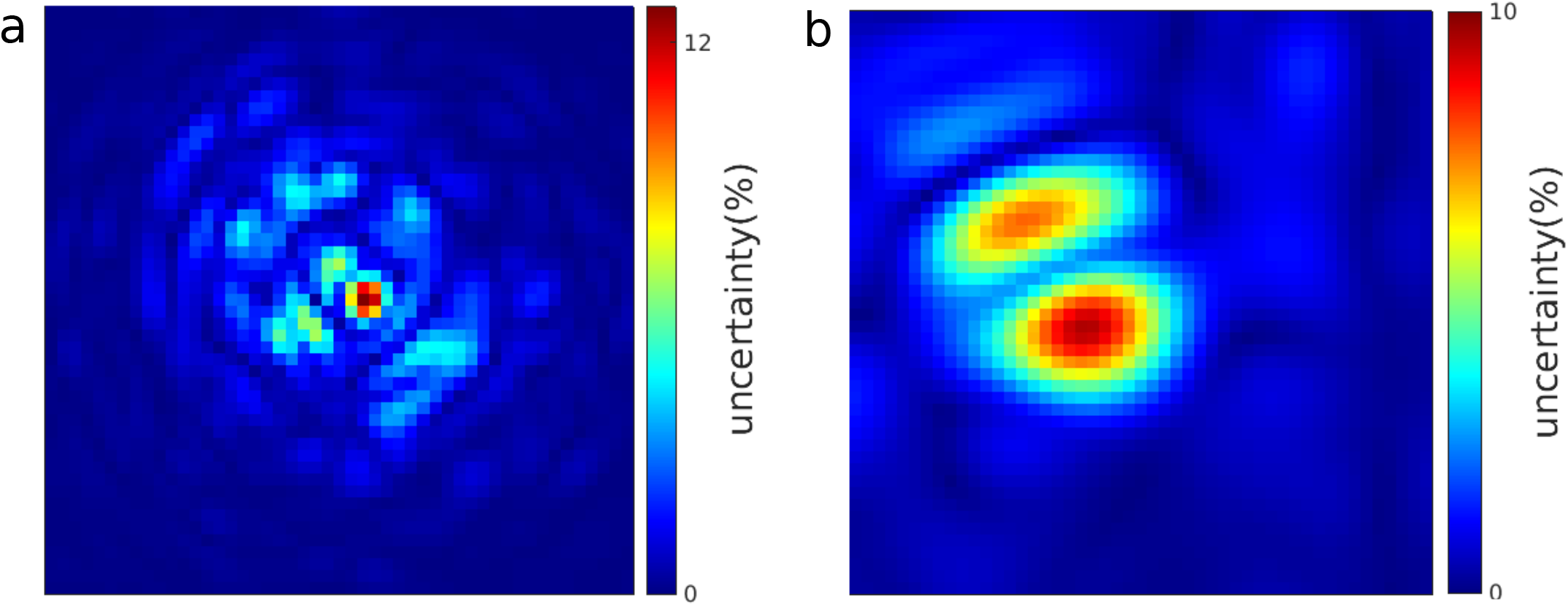
Pixel-wise uncertainty maps in PSF shape, at focal plane (Z=0), calculated as standard deviation divided by average of the last 2000 samples in the chain for every individual pixels. (a) uncertainty map corresponding to the PSF in Figs. 2-3 and Fig. S8c; (b) uncertainty map corresponding to the PSF in Fig. 5 and Fig. S8f.

**Fig. S11:**
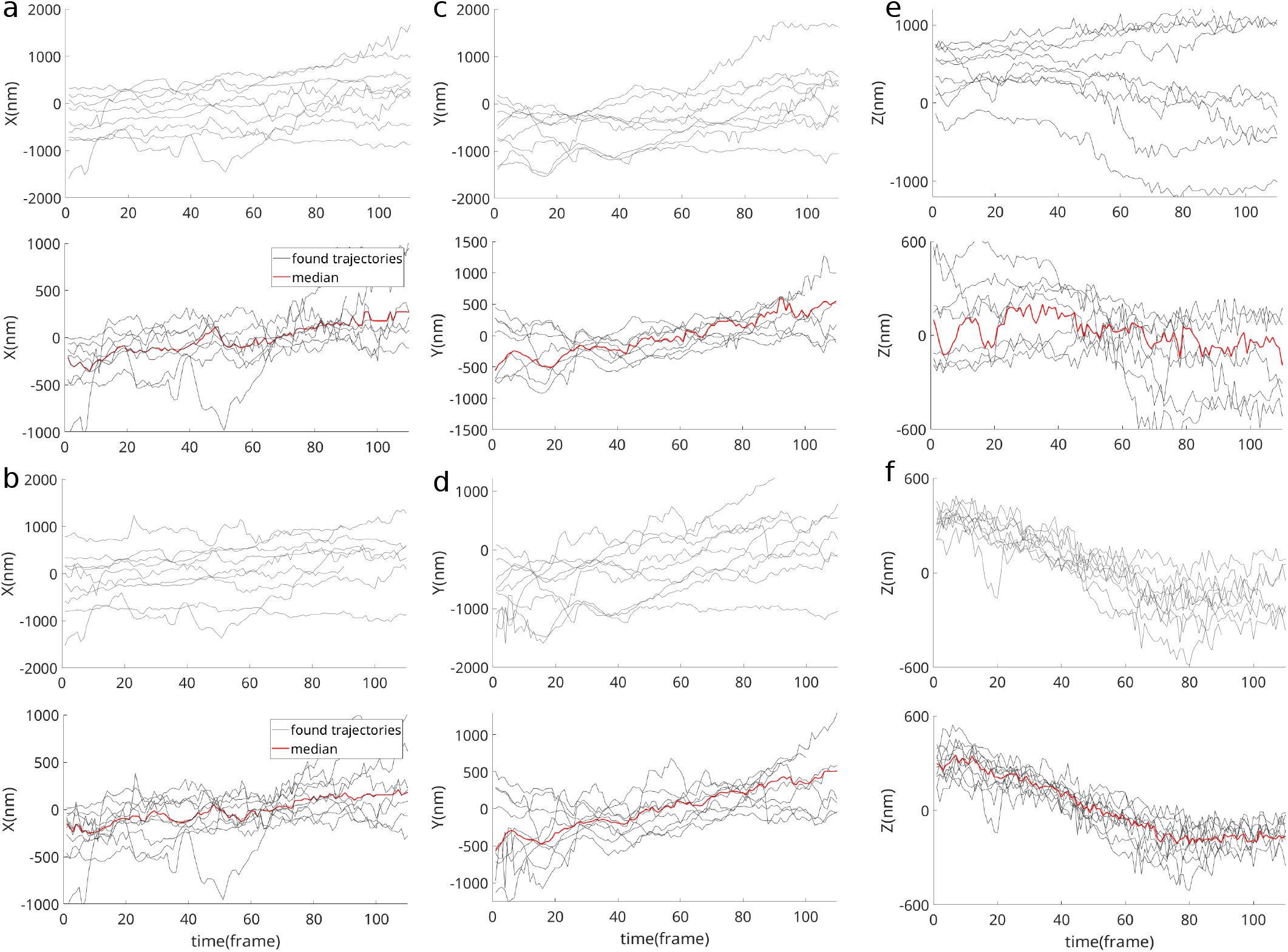
Tracking lytic granules in stimulated cells using Gaussian PSFs and SPT-PR. (a) The top panel shows trajectories (black) from 10 lytic granules in stimulated cells along the X-direction using Gaussian PSFs. In the bottom panel, for each lytic granule, the median position was subtracted from its trajectory before plotting. The red curve represents the median trajectory across lytic granules. (b) X-trajectories obtained using SPT-PR. (c) Y-trajectories obtained using a Gaussian PSF. (d) Y-trajectories obtained using SPT-PR. (e) Z-trajectories obtained using a Gaussian PSF. (f) Z-trajectories obtained using SPT-PR.

**Fig. S12:**
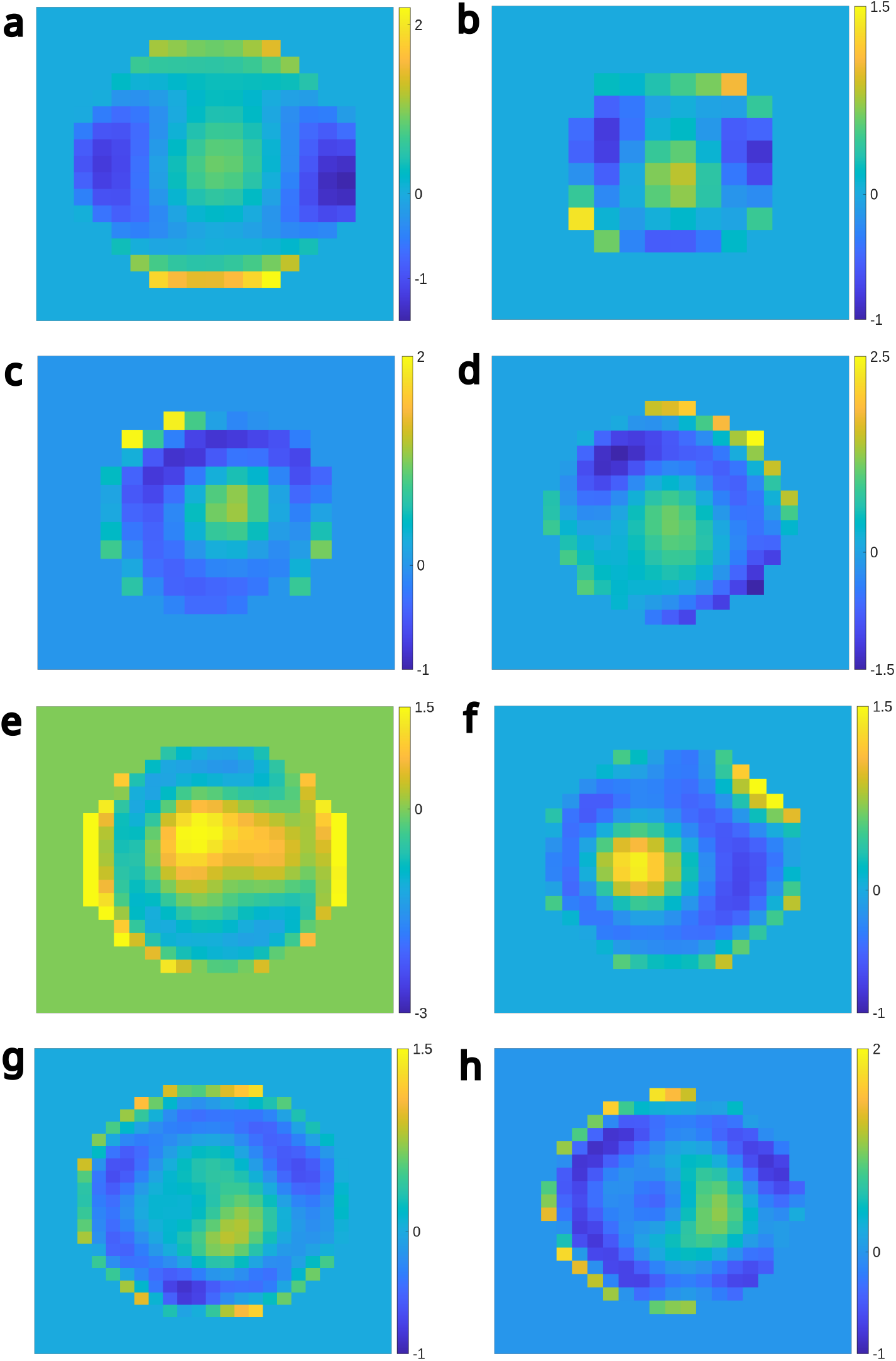
Pupil phases corresponding to the trajectories shown in Fig. 5 obtained by tracking lytic granules within (a-e) stimulated and (f-h) non-stimulated cells.

**Fig. S13:**
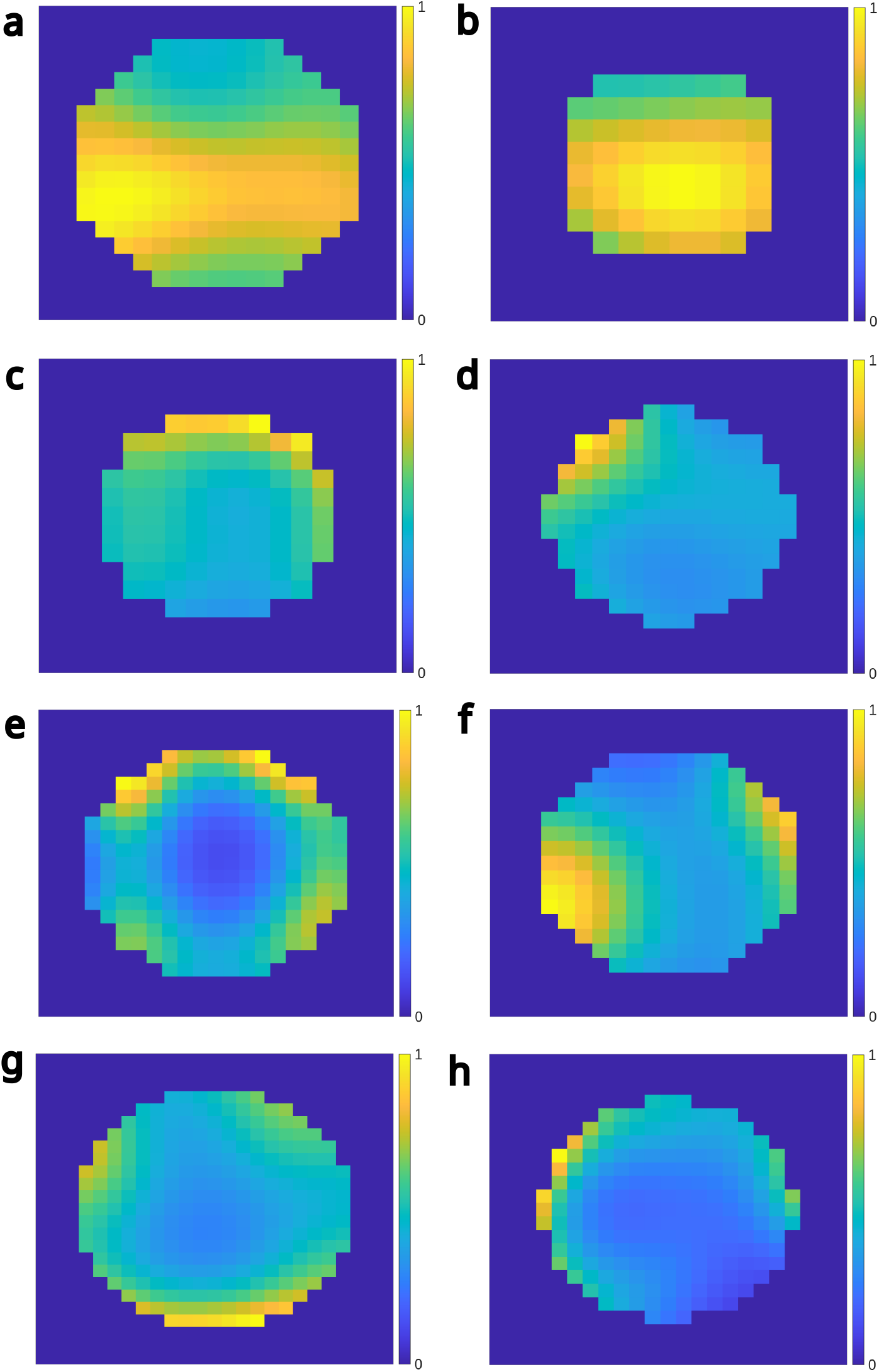
Pupil amplitudes corresponding to the trajectories shown in Fig. 5 obtained by tracking lytic granules within (a-e) stimulated and (f-h) non-stimulated cells.

### 1 Forward Model and Likelihood Derivation

In order to motivate eq. (1) from main text, here we start by the plane wave expansion of light’s electric field (Fourier representation) and employ it to describe light propagation [3,4]. To do so, we begin by a plane wave (single-frequency) given as

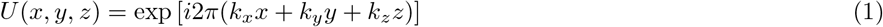

where *k*_*x*_,*k*_*y*_ and *k*_*z*_ are components of the vector 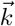 and 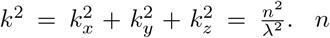 and λ are, respectively, the refractive index of the medium and the wavelength of light in vacuum. At the *z* = 0 plane, the plane wave is given by *U*(*x, y, z* = 0) = exp [*i*2*π*(*k*_*x*_*x* + *k*_*y*_*y*)]. Traveling along the z-axis, at *z* = *z*_0_ plane, the plane wave is then *U*(*x, y, z*_0_) = exp [*i*2*π*(*k*_*x*_*x* + *k*_*y*_*y* + *k*_*z*_*z*_0_)]. This can be expressed as follows

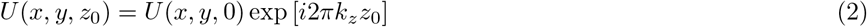

where exp [*i*2*πk*_*z*_*z*_0_] is single frequency plane wave propagator [3]

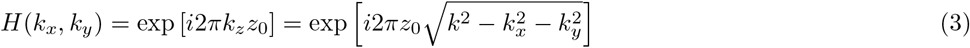

which is also called the defocus.

Now, an arbitrary field at *z* = 0 can be written as a superposition of single-frequency plane waves

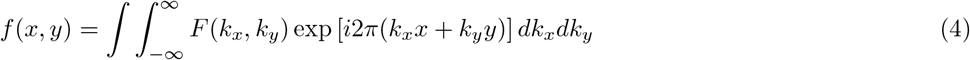

where *F*(*k*_*x*_, *k*_*y*_) is the Fourier transform of *f*(*x, y*). The transmitted field *f*(*x, y, z*_0_) is then

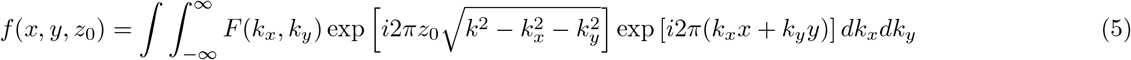

which is the Fourier transform of the product of *F*(*k*_*x*_, *k*_*y*_) and the defocus term.

In the case of microscopes, the Fourier transform of the light’s electric field is termed as the pupil function *𝒫* (*k*_*x*_, *k*_*y*_) which describes the wavefront and its distortions due to optical aberrations and/or extra phase intentionally added for point spread function (PSF) engineering. Here, Similar to eq. (5), we can write [4–6]

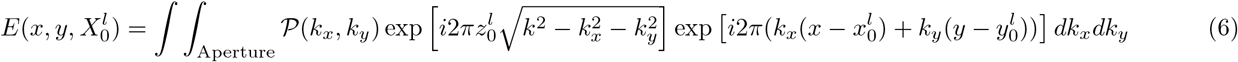

where the limits of the integral is also from zero up to the maximum spatial frequency transmitted by the objective/aperture, 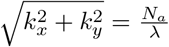, where *N*_*a*_ = *n* sin *θ* is the numerical aperture of the microscope. Here, *x* and *y* are the camera coordinates and the particle is located at 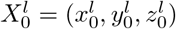 in the *l*th frame.

**Fig. S14:**
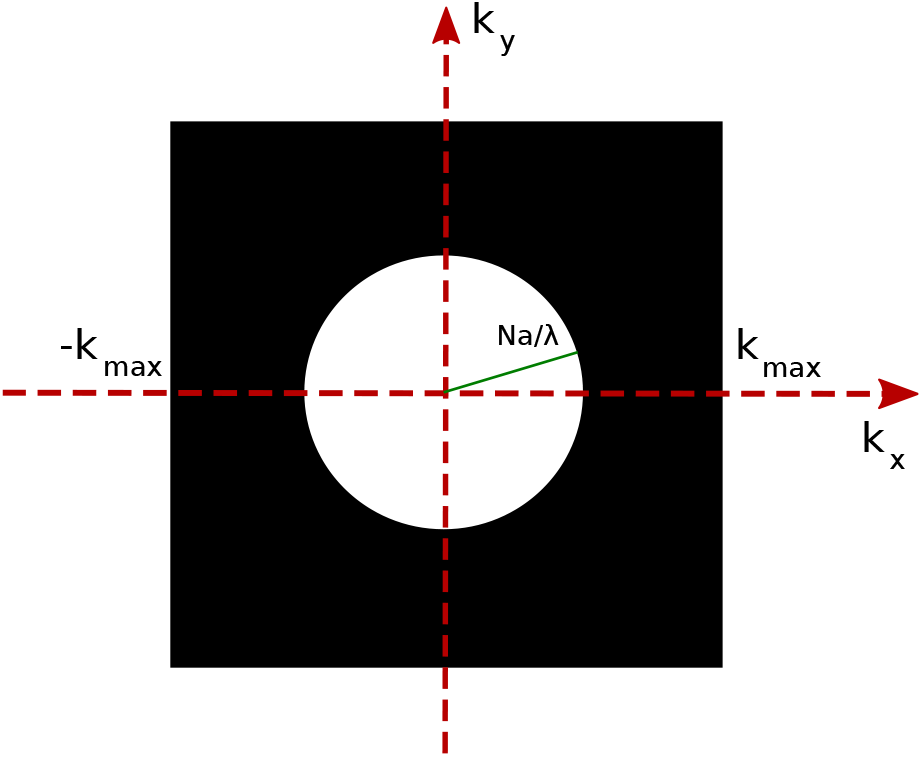
For convenience in performing the Fourier transformation in eq. (6), we define a mask in frequency domain ranging from −*k*_max_ to *k*_max_. The white circle represents the domain of frequencies passed by the microscope with a radius of *N*_*a*_/λ.

For convenience, we now define a mask, *M*(*k*_*x*_, *k*_*y*_), in the frequency domain where

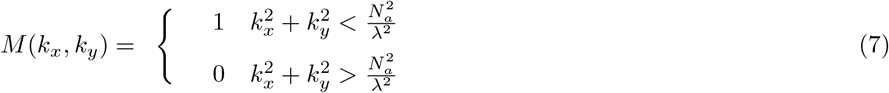

where we assume a limited size for the mask so that −*k*_max_ ≤ *k*_*x*_, *k*_*y*_ ≤ *k*_max_; see Fig. S14. Using the mask eq. (7), the integral in eq. (6) can be written as

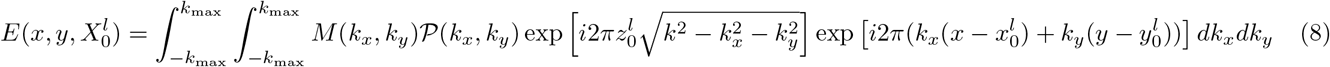

where the limits of the integral now coincide with the limits of the mask. The PSF is then the intensity of diffracted light, given by the square of the electric field

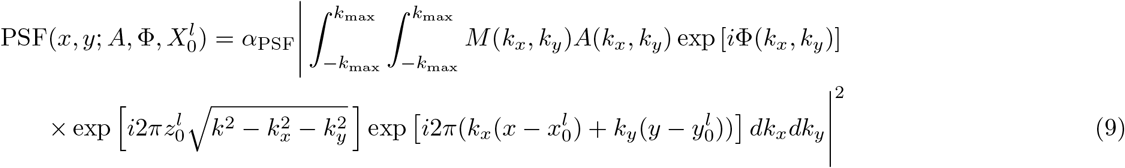

where *α*_PSF_ is the normalization constant and *𝒫* (*k*_*x*_, *k*_*y*_) = *A*(*k*_*x*_, *k*_*y*_) exp [*i*Φ(*k*_*x*_, *k*_*y*_)]. Here, *A* ((*k*_*x*_, *k*_*y*_) and Φ(*k*_*x*_, *k*_*y*_) are, respectively, amplitude and phase of the pupil function and are real quantities. Using the above PSF, the expected photon counts over a pixel over a single exposure time (frame) is given by

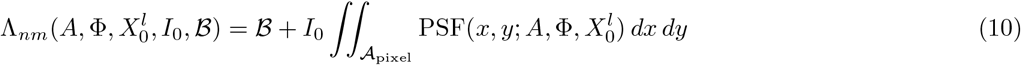

where *ℬ, I*_0_ and *𝒜*_pixel_ are, respectively, the background photon count per pixel, total photon count from the particle and the pixel area [6–8]. Eq. (10) can be easily extended to the case with multiple particles by summing over the photon contribution from each particle

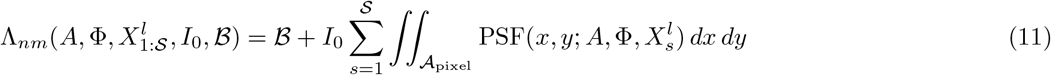

where *s* counts the *𝒮* particles present within the model and 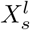 is the *s*th particle location at time point *l*.

Although we have so far discussed an imaging model using continuous coordinates, calculation of integrals in eq. (9) and eq. (10) are often computationally expensive. As such, it is convenient to approximate these integrals by sums. To do so, we first note that the PSF is typically measured over a region of interest (ROI) with *N* × *N* pixels with pixel size of *a*. Further, we presume *n, m* = 1, …, *N*, respectively, count rows and columns of pixels in that ROI. The Fourier transform of the PSF is therefore also given over a ROI of *N* × *N* pixels with pixel size of 1*/*(*aN*), *i*.*e*., grid with grid step size of 1*/*(*aN*), in the frequency domain, where the associated discrete frequencies along *x* and *y* axes are given by *k*_*µ*_, 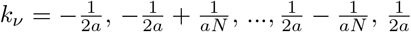. The integral in eq. (9) can thus be written as a sum

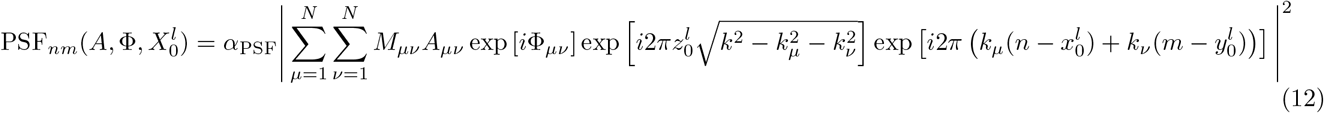

where 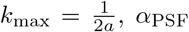 is the normalization constant and *µ, ν* count pixels (samples) in the Fourier domain. To further reduce the computational cost, the above sum can be calculated using fast Fourier transform (FFT)

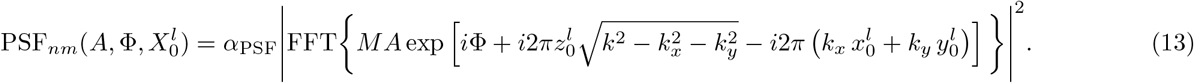

where *M, A* and Φ represent the entire se(ts of {*M*_*µν*_, *A*_*µν*_, Φ_*µν*_}. The PSF model presented in eq. 13 often re(sults in a slightly sharper PSF than the measured PSF as observed in Refs. [5, 9–11]. This effect can be modeled by convolving the PSF given by eq. 13 with a Normal distribution as follows [5, 9–11]

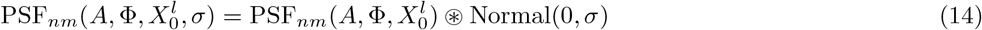

where 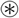 denotes convolution.

The above equation yields the PSF values at the pixel centers which can be multiplied by the pixel area to approximate the PSF integral required in calculating the expected photon counts over pixels; see eq. (10). However, particle localization based on such approximation would not provide sufficient precision and accuracy [12] for sub-diffraction-limited tracking [1]. To remedy this issue, we need to calculate pixel values with resolutions below the data pixel size and sum over the obtained values to gain a more accurate approximation of the integral in eq. (10). To calculate PSF values below the pixel size, it is common to use zero-padding in Fourier domain, which basically means increasing the sizes of *M, A* and Φ in eq. (13) to larger frequencies by adding zeros to them [13]. Although the zero-padding method does not yield any additional information, it gives PSF values below the pixel size via interpolation.

Now, assuming that the zero-padding results in *J* samples of PSF per pixel, we show the result by 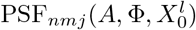. Therefore, the expected photon count at the *n*th and *m*th pixel over the *l*’th frame is given by (see eq. (10))

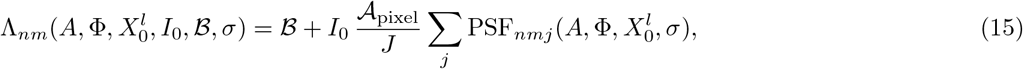

where 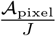 is the area of the pixel divided into *J* sub-pixels. Now, we can construct a likelihood employing the obtained expected photon counts and use it to track the particle across *L* frames. However, due to the symmetry of the PSF with respect to the focal plane the localization accuracy along the axial direction would be insufficient. One way to address this issue is multi-plane imaging resulting in multiple simultaneous slices of the PSF along the axial direction with known separation. The expected photon counts for each plane can be computed similar to eq. (15)

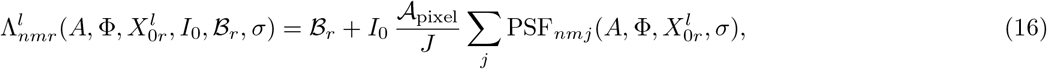

where *r* counts *R* planes in the imaging setup. Therefore, assuming photon shot noise and the camera noise, the total likelihood is [4, 14]

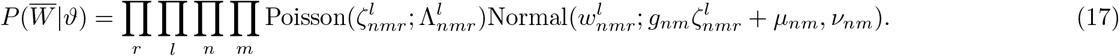

where 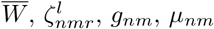 and *ν*_*nm*_ are, respectively, the measured photon counts over the entire sequence across all frames and all focal planes, observed photon counts, camera gain, offset and noise variance. However, this likelihood has a complicated form and leads to intense computational complexity. Therefore, to reduce the computational complexity, we used the following approximation [4, 14]

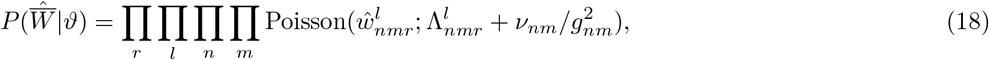

where 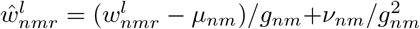, and 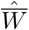 represents the entire set of 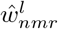 . Moreover, here 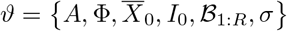 shows the entire set of unknowns and we use 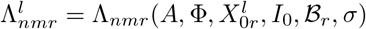 for succinctness hereafter.

Here a note is warranted on the multi-plane setup. In such setpus, the axial locations are often registered to be the same across all the planes and the interplane distance along the axial direction (*z*) is known. As such, one of the planes is assumed to be the reference plane and the particle location is only learned in that plane. The particle location in the remaining planes are deterministically related to the location in the reference plane using translation and rotation which can be simultaneously performed using affine transformations [15]. The respective shift and rotation across the planes can be found using calibration data from bright beads. Here, we assume that the reference plane is *r* = 1 and only calculate particle location in this plane, namely 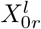.

#### 1.1 Freely diffusing particle

So far, we have described the imaging model. Here, we assume a diffusing particle that freely diffuses and explore the area and therefore its location changes over time as follows

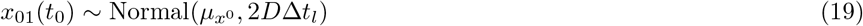

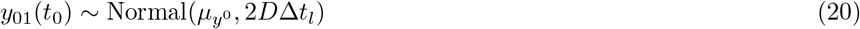

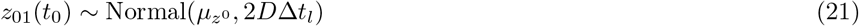

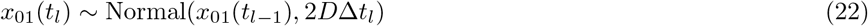

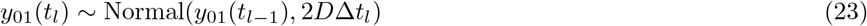

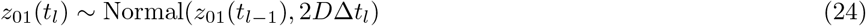

where *l* counts frames and Δ*t*_*l*_ = *t*_*l*_ − *t*_*l*−1_ is the exposure time for the *l*th frame, and *D* is the diffusion coefficient. *x*_01_(*t*_0_), *y*_01_(*t*_0_) and *z*_01_(*t*_0_) are the particle’s initial location in the reference plane assumed to be at a random position across the ROI taken from normal distributions in eq. (19)-(21). As mentioned before, the particle’s position in the second plane is deterministically related to the first (reference) plane by an affine transformation. Finally, using the generated sequence of particle locations, we can compute the expected photon counts 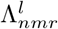 given by eq. (15) and use it to find the likelihood in eq. (18).

**Fig. S15:**
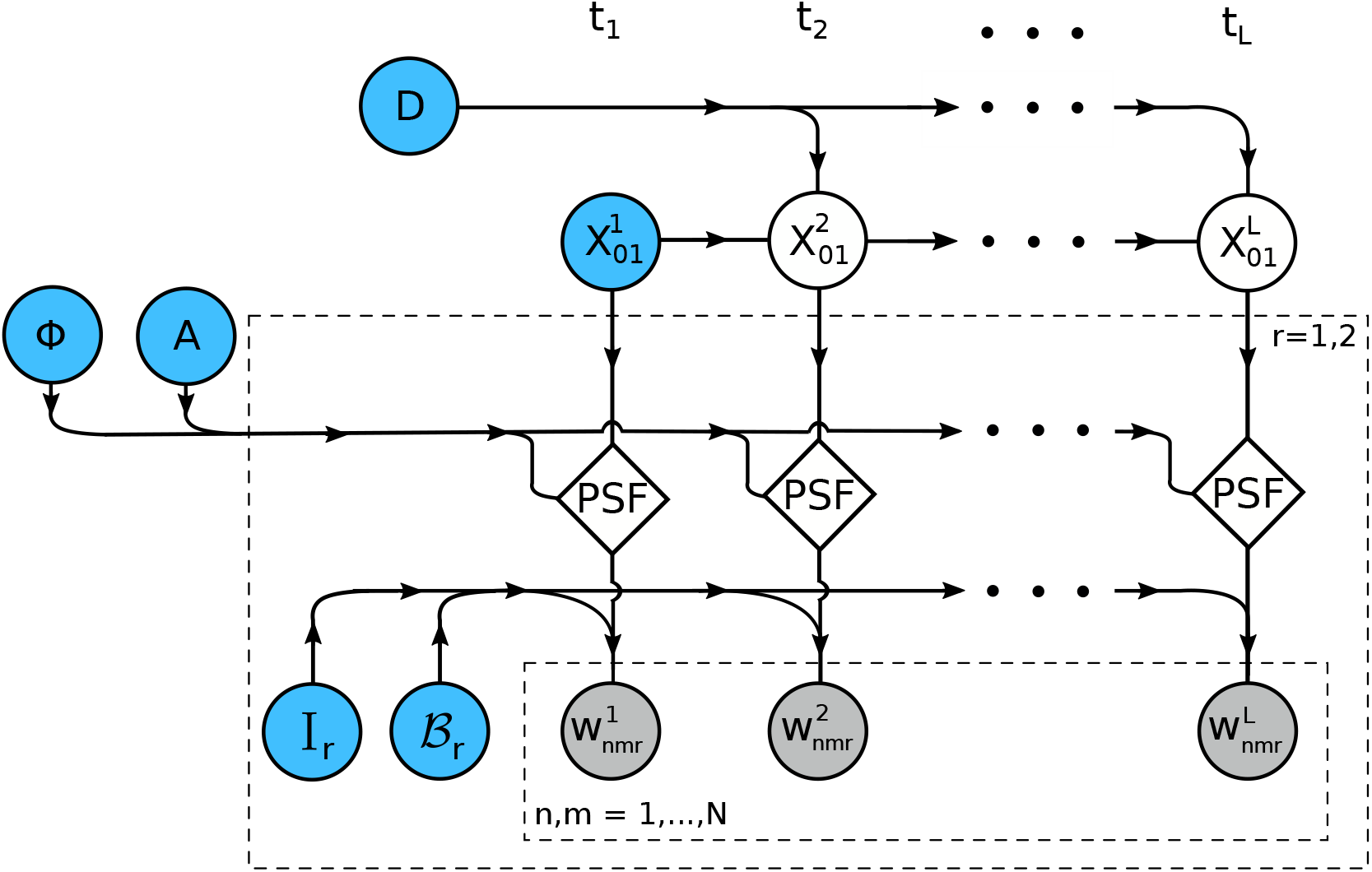
Graphical Model depicts hierarchical dependencies among parameters. This graphical model assumes a single particle; 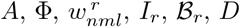 and 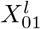, respectively, stand for the amplitude and phase of pupil function, data frame images, photon emission rate and background emission rate at the r-plane the diffusion constant, and particle’s location in the first (reference) plane. *l* = 0, …, *L, r, n* and *m* count the data acquisition times, the imaging planes, pixels row and column indices, respectively. The blue, gray and blank circles, respectively, represent unknown parameters, hidden parameters and data. The diamonds represent variables that can be deterministically calculated from other parameters. Note that the parameter *D* is not directly connected to the data, indicating its second order hierarchy.

### 2 Inverse Model

Here, we first give a summary of the inverse model including priors used on unknowns and the likelihood model and then provide details in what follows.

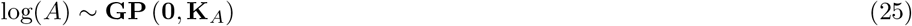

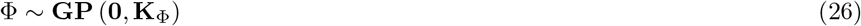

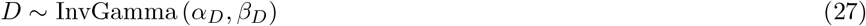

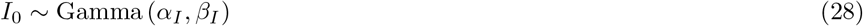

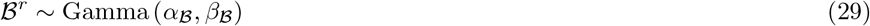

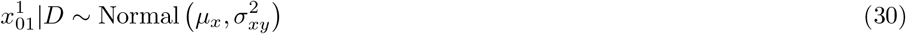

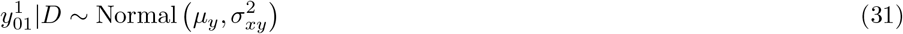

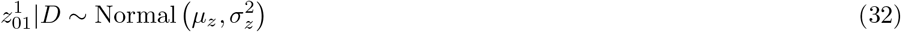

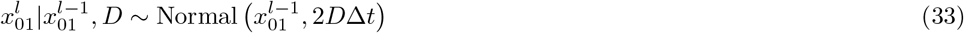

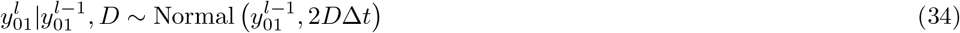

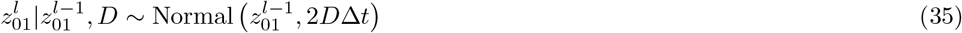

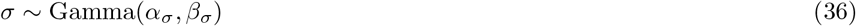

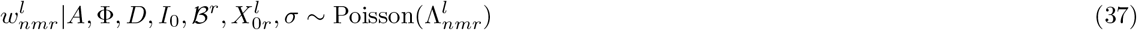

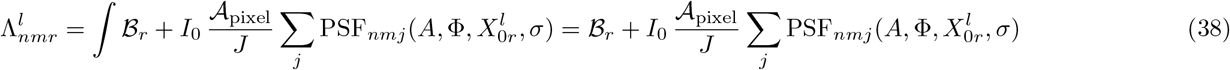

where ∼ indicates that the parameter on the left follows the distribution given on the right hand side. Moreover, **K**_Φ_ and **K**_*A*_ are the covariance kernels for Gaussian process (GP) priors given by

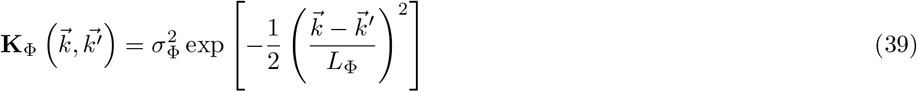

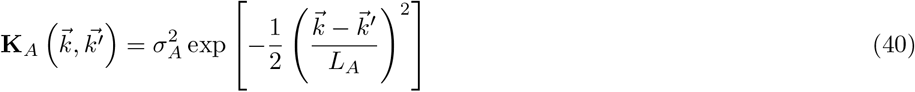

where 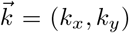 are the coordinates in the Fourier domain and *σ*_Φ_, *σ*_*A*_, *L*_Φ_ and *L*_*A*_ are positive quantities used to adjust correlations across space. It is assumed that the PSF is already integrated over space.

#### 2.1 Making Inference about

The target distribution of the phase of the pupil function, Φ, is given by

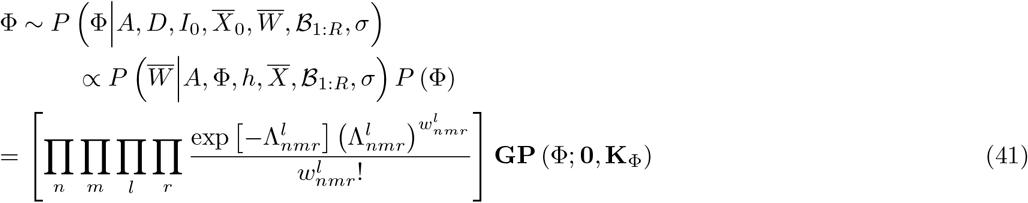

We use the Gaussian processes technique to sample the pupil phase (Φ). To do so, the pupil phase is sampled at a mesh grid of test points with the same number of elements as the data ROIs following the same procedure as Ref. [16]. We then subtract contributions of the first four Zernike polynomials, *i*.*e*., offset, *x*-tilt, *y*-tilt and defocus, where the last three terms are associated to shifts in particle’s location [4], from the sampled phase. These contributions are given as *α*_*s*_*Ƶ*_*s*_ for the *s*-th Zernike polynomial with

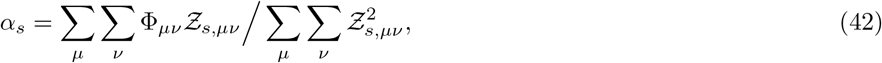

where *Ƶ*_*s,µν*_ is the discretized *s*-th Zernike polynomials. *µ* and *ν* count discrete elements of the Zernike polynomial along the first and second dimensions; see eq. (12). The result is thus the phase due to only optical aberrations and is employed in the subsequent calculations. Since the likelihood is not conjugate to the GP prior, we use Metropolis-Hastings (MH) technique to sample the posterior (41)

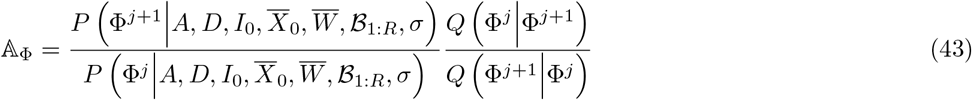

where *Q* is the proposal distribution and *j* count samples. We use the GP prior itself as the sampling distribution and therefore the acceptance ratio is given by the likelihood ratios

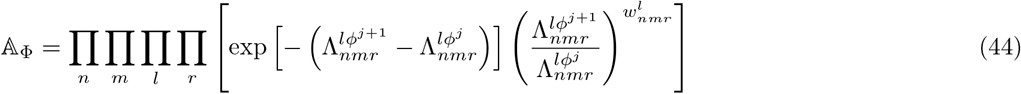

where 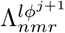 is calculated using the proposed phase *ϕ*^*j*+1^.

#### 2.2 Making Inference about *A*

The target distribution of the pupil function amplitude, *A*, is given by

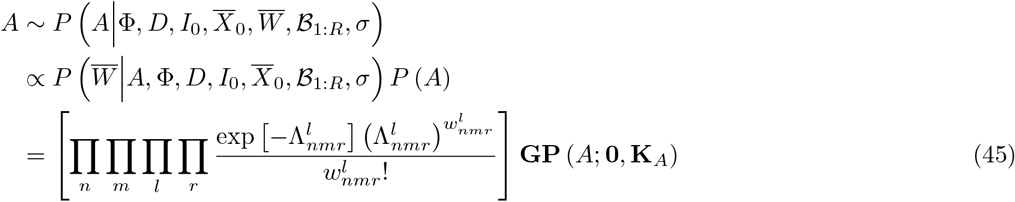

We use Gaussian process technique to sample the amplitude of the pupil function. However, while the amplitude is a positive quantity, GPs allow negative values as well. Thus we make a substitution *A* = *e*^*χ*^ and learn *χ* which can be either negative or positive. Similar to Φ, we use MH technique to sample the posterior (45) and we also use the Gaussian process prior as proposal distribution. Therefore the acceptance ratio is given by

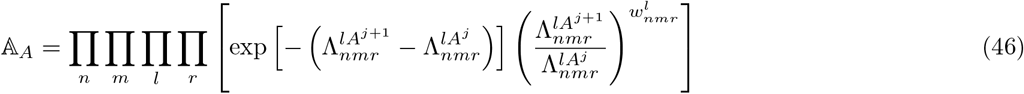

where 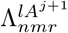 is calculated using the proposed amplitude.

#### 2.3 Making Inference about D

The target distribution of the diffusion coefficient, *D*, is given by

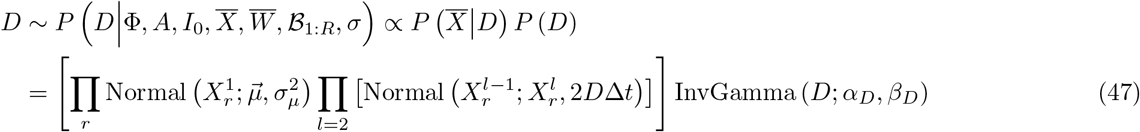

The InvGamma prior is conjugate to the likelihood and the target posterior has a closed form given by

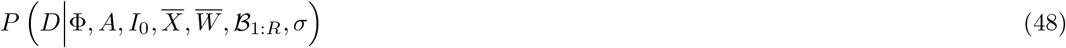

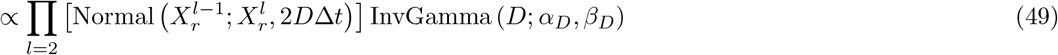

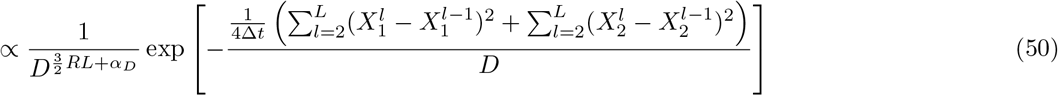

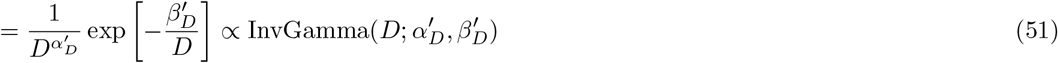

where we dropped the prefactors that do not depend on *D* in the last step. Furthermore, here, we have

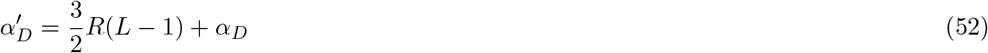

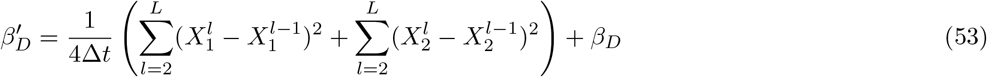

where *R* is the number of planes. Therefore, the diffusion constant, *D*, can be directly sampled form the posterior eq. (51).

#### 2.4 Making Inference about *I*_0_

The target distribution of *I*_0_ is given by

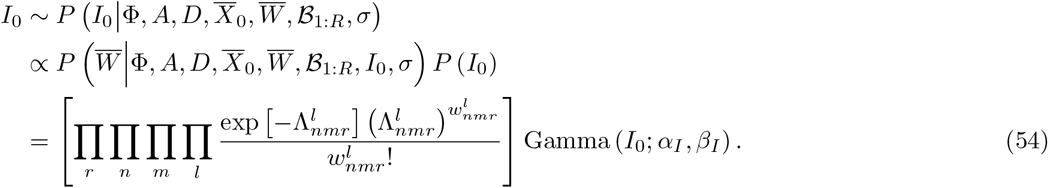

Since the posterior eq. (54) does not have a closed form, we use MH technique to draw samples

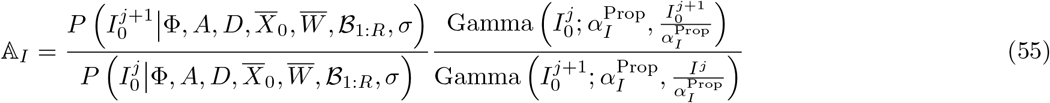

where we used the gamma distribution as proposal distribution

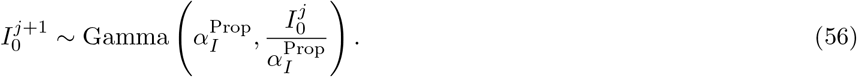

#### 2.5 Making Inference about *ℬ*_*r*_

The target distribution of *ℬ*_*r*_ is given by

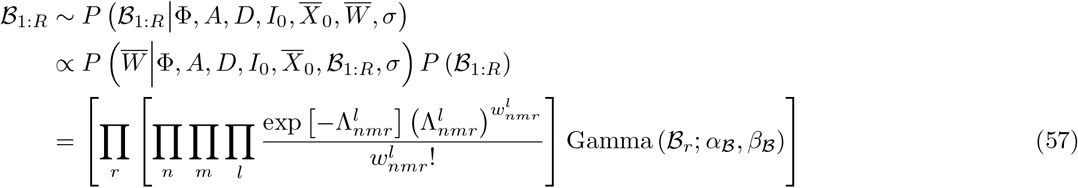

Since the posterior eq. (57) does not have a closed form, we use MH technique to draw samples

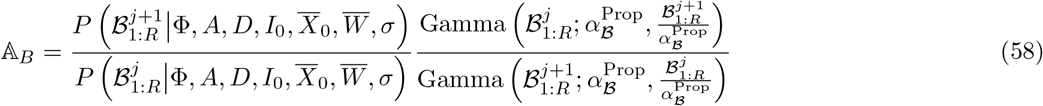

where we used the gamma distribution as proposal distribution

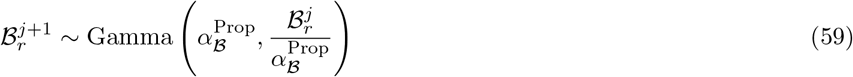

#### 2.6 Making Inference about 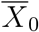

The target distribution for 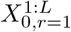 is given by

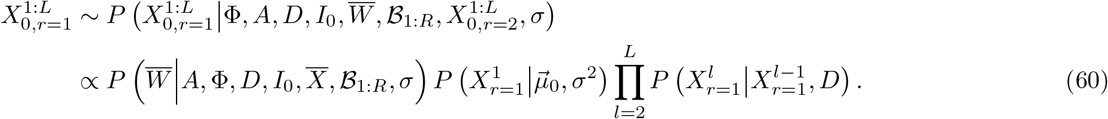

Here, we sample particle’s trajectory in the reference plane, *i*.*e*., *r* = 1, and locations in other planes can be deterministically calculated. The Z-trajectory is initialized to all zero positions, and the initial X and Y trajectories are set to the coordinates of the brightest pixel in each frame.

We propose new trajectories by:

1. selecting a random frame and modifying the corresponding location:

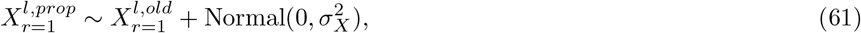
2. modify the particle locations across all frames using hit-and-run procedure [17]

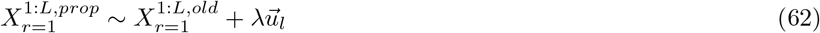

where 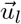 and λ are, respectively, a unit vector with a random direction in space and the amplitude of the move in that direction. Note that the amplitude of the move in all directions are similar. The acceptance ratio of both jumps can be calculated as follows

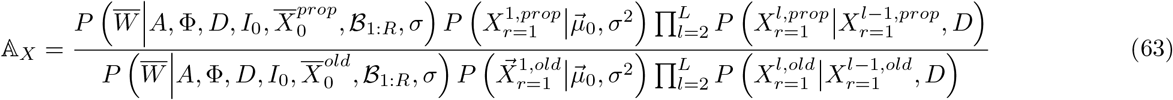

where the proposal distributions are canceled.

#### 2.7 Making Inference about about *σ*

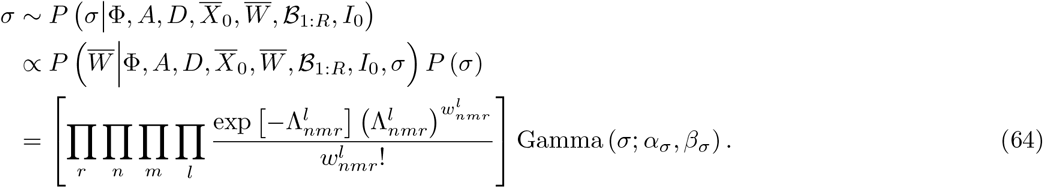

Since the posterior eq. (64) does not have a closed form, we use MH technique to draw samples

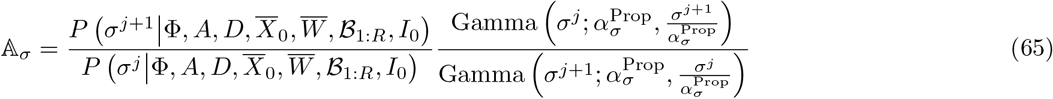

where we used a gamma distribution as the proposal distribution

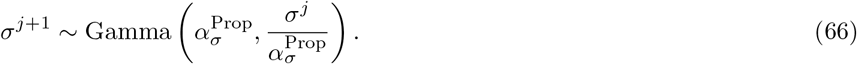

### 3 Direct Calculation of Diffusion Coefficient Using a Given Trajectory

Using a given trajectory segment 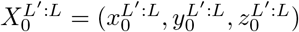 from time point *L*^∓^ to *L*, the corresponding diffusion coefficient can be found directly assuming a Brownian motion. The likelihood for the Brownian motion is given as

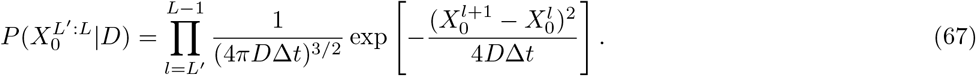

Now, we take the derivative of the log-likelihood with respect to *D*

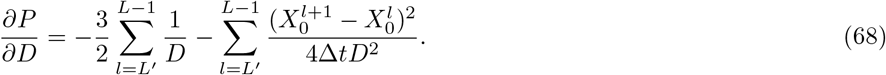

Setting this to zero and solving for *D*, we obtain

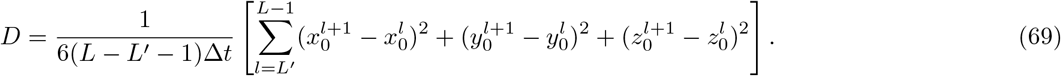

